# De-novo purine biosynthesis is a major driver of chemoresistance in glioblastoma

**DOI:** 10.1101/2020.03.13.991125

**Authors:** Jack M Shireman, Fatemeh Atashi, Gina Lee, Eunus S. Ali, Miranda R. Saathoff, Cheol H. Park, Shivani Baisiwala, Jason Miska, Maciej S. Lesniak, James C. David, Roger Stupp, Priya Kumthekar, Craig M. Horbinski, Issam Ben-Sahra, Atique U. Ahmed

**Affiliations:** Department of Neurological Surgery, Department of Biochemistry and Molecular Genetics, Feinberg School of Medicine, Northwestern University, Chicago, IL USA 60616

## Abstract

This year nearly 20,000 lives will be lost to Glioblastoma (GBM), a treatment-resistant primary brain cancer. In this study, we identified a molecular circuit driven by epigenetic regulation that regulates the expression of ciliary protein ALR13B. We also demonstrated that ARL13B subsequently interacts with purine biosynthetic enzyme IMPDH2. Removal of ARL13B enhanced TMZ-induced DNA damage by reducing de-novo purine biosynthesis and forcing GBM cells to rely on the purine salvage pathway. Furthermore, targeting can be achieved by using an FDA-approved drug, Mycophenolate Moefitil. Our results suggest a clinical evaluation of MMF in combination with TMZ treatment in glioma patients.

## Introduction

Glioblastoma (GBM) is an almost universally lethal primary brain tumor that will take the lives of roughly 20,000 people this year alone^1^. Despite an aggressive standard of care therapy, the median survival of a patient with GBM is just 20 months^2^. This unfavorable prognosis is mainly due to the high rate of recurrence, as recurrent GBMs are often unresponsive to all avenues of therapy. Although it has become a significant focus of recent research, the true mechanistic underpinnings of resistance to therapies are yet to be elucidated. The heterogeneous nature of GBM tumors combined with the existence of resistant subpopulation, such as tumor-initiating cells (TIC), are considered to be significant drivers of resistance^3–5^.

TIC’s are known for their adaptive or “plastic” nature, which allows for the acquisition of stemness in response to appropriate microenvironmental cues such as hypoxia or therapeutic stress ^6–9^. Research has shown that such plasticity is regulated by changes in permissive epigenetic states, which allow cells to employ rapid context-dependent regulation of genes that may be necessary for conferring fitness during therapy^10–12^. Unfortunately, therapeutically actionable targets whose inhibition would limit TIC adaptation have proven challenging to uncover.

A canonical driver of context-dependent epigenetic plasticity is the Polycomb Repressor Complex (PRC) and its catalytic subunit Enhancer of Zeste Homologue 2 (EZH2). The PRC2/EZH2 complex not only required for different neurodevelopmental processes but also associated with disease pathogenesis including GBM^13–20^. Tumor cells can also adapt to treatment metabolically by rapidly altering core metabolic pathway activity^21, 22^. In this study, we show that resistance to temozolomide (TMZ), as part of the standard of care for newly diagnosed GBM, is associated with EZH2/PRC2 regulated ARL13B. Further, we demonstrated an interaction between ARL13B and IMPDH2, a rate-limiting enzyme of purine biosynthesis, that impacts GBM’s adaptive response to TMZ by inhibiting purine salvaging. Disruption of IMPDH2 activity by using an FDA approved compound Mycophenolate Moefitil (MMF) significantly increased the therapeutic efficacy of TMZ. MMF extends the survival of patient-derived xenograft (PDX) models of mice across all GBM subtypes. Our study, therefore, provides evidence of a rapidly clinically translatable opportunity to enhance the efficacy of alkylating agents in GBM.

## Materials and Methods

### Animal studies

Athymic nude mice (NU(NCr)-Foxn1nu; Charles River Laboratory) were maintained according to all Institutional Animal Care and Use Committee guidelines. In compliance with all applicable federal and state statutes governing animal use in biomedical research, the mice were housed before and during the study in a temperature- and humidity-controlled room following a strict 12-hour light/dark cycle.

### Cell culture

Patient-derived xenograft (PDX) glioblastoma specimens GBM5, GBM6, GBM43, and GBM52, were obtained from Dr. C. David James (Northwestern University) and maintained for in vitro experiments in DMEM (Thermo Fischer Scientific) supplemented with 1% fetal bovine serum (FBS; Atlanta Biologicals) and 1% Antibiotic Antimycotic Solution (Corning) according to established protocols with slight alterations^23^. For the generation of shRNA knockdown lines, lentivirus particles were made using HEK293 cells (ATCC) transfected with 2nd generation packaging/envelope plasmids (Dr. Yasuhiro Ikeda, Mayo Clinic) and shRNA clones (GeneCopoeia). U251 cells were obtained from American Type Cell Culture and maintained for in vitro experiments in DMEM supplemented with 10% FBS and 1% Antibiotic Antimycotic Solution. CRISPR knockout of U251 cells was created by direct transfection with Cas9 nuclease and sgRNA targeting ARL13B (Dharmacon). All cells were passaged by washing one time with phosphate-buffered saline solution (PBS; Gibco) and detached using 0.25% trypsin/2.21mM EDTA (Corning).

### Flow cytometry

Flow cytometric analysis was performed on homogenized tumor tissue after HLA-based isolation from murine brains and adherent PDX cells after trypsinization. Cells were washed three times in PBS, incubated in fixation buffer (eBioscience) for 30 minutes, and permeabilization buffer for 30 minutes at room temperature. Cells were washed and incubated in primary conjugated antibodies (EZH2, BD Bioscience; CD133, Miltenyi; CD15, Biolegend; Sox2, Sino Biological Inc.; HIF1a, Biolegend; HIF2a, Abcam; and ENT1, Abcam) for 30 minutes at room temperature. Cells were washed a final time, resuspended in 0.5% BSA + TritonX (Sigma) FACS buffer, and analyzed using the BD LSRFortessa 6-Laser (BD Biosciences).

### Western blot

After trypsinization, PDX cells were lysed using mammalian protein extraction reagent (mPER; Thermo Scientific) buffer, and protein concentration were quantified via Bradford Assay. Protein samples were prepared by combining with reducing SDS (Alfa Aesar) and boiling at 95°C for 10 minutes. Following denaturation, samples were loaded into 10% polyacrylamide gels, transferred to nitrocellulose membrane (Bio-Rad), and blocked in TBS-T/5% milk solution. Membranes were incubated overnight in primary antibodies (EZH2; Cell Signalling; Beta Actin, ProteinTech; ARL13B, ProteinTech; IMPDH2, Abcam; Oct4, Cell Signaling; Sox2, Cell Signaling; Shh, Cell Signaling; Sufu, Cell Signaling; IFT43, ProteinTech; and ENT1, Abcam) at 4°C and for 2 hours at room temperature in HRP-conjugated secondary antibodies (Cell Signaling).

### Immunoprecipitation

Following trypsinization, cells were lysed using mPER buffer, and protein concentration were assessed via Bradford Assay. Samples were incubated overnight at 4°C for antibody crosslinking (anti-HA, ProteinTech; Normal Rabbit IgG, Cell Signalling; ARL13B, ProteinTech; and IMPDH2, Abcam) at a dilution of 1:66. The following morning Protein A/G UltraLink Resin (Thermo) was added to each reaction at a dilution of 1:4 and incubated at room temperature for 2 hours. Precipitates were washed three times with 500uL of mPER buffer and pelleted in a tabletop microfuge for 5 minutes at 2500 xg. Antibody-antigen complexes were eluted in 2X, reducing SDS at 55°C for 10 minutes. Finally, the samples were boiled at 95°C for 5 minutes.

### Immunofluorescence

For frozen, OCT-embedded (Sakura) tissue, sections were thawed in a 37°C humid chamber for 20 minutes and fixed in 4% paraformaldehyde (PFA; Thermo) for 15 minutes at room temperature. For adherent cells, 8-well chamber slides were taken from 37°C incubators following completion of treatments and fixed 4% PFA for 10 minutes at room temperature. Following fixation, slides were washed two times in 0.05% Tween/PBS solution (PBS-T, Fischer Bioreagents) for 5 minutes and blocked in 10% BSA/0.3% Triton-X/PBS solution at room temperature for 2 hours. Samples were incubated overnight at 4°C in primary antibody mixtures (HIF1α, Biolegend; EZH2, BD Bioscience; ARL13B, ProteinTech; IMPDH2, Abcam; SMO, Santa Cruz; and Gli1, R&D Systems) diluted with 1% BSA/0.3% Triton-X/PBS solution. For unconjugated primaries, the following morning slides were washed three times and incubated at room temperature for 2 hours in secondary antibody mixtures (Invitrogen). Finally, slides were washed three times and mounted using ProLong™ Gold Antifade reagent with Dapi (Invitrogen). Slides were imaged using a fluorescence microscope (Model DMi8; Leica).

### Microarray

Following trypsinization, RNA was extracted from cells using the RNeasy Plus Mini Kit (Qiagen). Microarray analysis was performed using 1,000ng of RNA per manufacturer’s guidelines (Illumina). HumanHT12 (48,000 probes, RefSeq plus EST) was used for each microarray. All microarrays were performed in triplicate.

### Mass spectroscopy

Samples were run on an SDS-PAGE gel, and a gel band was subject to in-gel digestion. Gel band was washed in 100 mM Ammonium Bicarbonate (AmBic)/Acetonitrile (ACN) and reduced with ten mM dithiothreitol at 50°C for 30 minutes. Cysteines were alkylated with100 mM iodoacetamide in the dark for 30 minutes at room temperature. Gel band was washed in 100mM AmBic/ACN before adding 600 ng trypsin for overnight incubation at 37 °C. The supernatant containing peptides was saved into a new tube. The gel was washed at room temperature for ten minutes with gentle shaking in 50% ACN/5% FA, and the supernatant was saved to peptide solution. Washing was repeated each by 80% ACN/5% FA, and 100% ACN, and all supernatant was saved into a peptide solution then subject to speed vac drying. After lyophilization, peptides were reconstituted with 5% ACN/0.1% FA in water and injected onto a trap column (150 µm ID X 3cm in-house packed with ReproSil C18, 3 µm) coupled with an analytical column (75 µm ID X 10.5 cm, PicoChip column packed with ReproSil C18, 3 µm) (New Objectives, Inc., Woburn, MA). Samples were separated using a linear gradient of solvent A (0.1% formic acid in water) and solvent B (0.1% formic acid in ACN) over 120 minutes using a Dionex UltiMate 3000 Rapid Separation nanoLC (ThermoFisher Scientific). MS data were obtained on an Orbitrap Elite Mass Spectrometer (Thermo Fisher Scientific Inc, San Jose, CA). Data were analyzed using Mascot (Matrix Science, Boston, MA) v.2.5.1 against the Swiss-Prot Human database (2019), and results were reported at 1% FDR in Scaffold v.4.8.4 (Proteome Software, Portland, OR). For stable isotope (13C5-hypoxanthine) labeling experiments, dried pellets were resuspended using 20 ul LC-MS grade water for mass spectrometry. 10 µL were injected and analyzed using a 5500 QTRAP triple quadrupole mass spectrometer (AB/SCIEX) coupled to a Prominence UFLC HPLC system (Shimadzu) via selected reaction monitoring (SRM) ^24^. Some metabolites were targeted in both positive and negative ion modes for a total of 287 SRM transitions using pos/neg polarity switching. ESI voltage was +4900V in positive ion mode and –4500V in negative ion mode. The dwell time was 3 ms per SRM transition, and the total cycle time was 1.55 seconds. Approximately 10-14 data points were acquired per detected metabolite. Samples were delivered to the MS via normal phase chromatography using a 4.6 mm i.d x 10 cm Amide Xbridge HILIC column (Waters Corp.) at 350 µL/min. Gradients were run starting from 85% buffer B (HPLC grade acetonitrile) to 42% B from 0-5 minutes; 42% B to 0% B from 5-16 minutes; 0% B was held from 16-24 minutes; 0% B to 85% B from 24-25 minutes; 85% B was held for 7 minutes to re-equilibrate the column. Buffer A was comprised of 20 mM ammonium hydroxide/20 mM ammonium acetate (pH=9.0) in 95:5 water: acetonitrile. Peak areas from the total ion current for each metabolite SRM transition were integrated using MultiQuant v2.0 software (AB/SCIEX). For stable isotope labeling experiments, custom SRMs were created for expected 13C incorporation in various forms for targeted LC-MS/MS.

For all extractions, the remaining pellets were resuspended in 8 M Urea /10 mM Tris, pH 8, heated at 60°C degrees for 30 min with shaking, centrifuged, and protein concentration in the supernatant was quantified using Bradford kit. Peak areas of metabolites detected by mass spectrometry were normalized to the median and then normalized to protein concentrations.

### Immunohistochemistry

Slides were placed in an oven at 60°C for 1 hour; then paraffin was removed using Xylene (3×5min), 100% ETOH (2×3 min), and 95% ETOH for 3 mins. Slides were then placed into a retrieval solution (ph. 6) and incubated in a Biocare Medical Decloaking Chamber set to 110°C for 5 minutes. After cooling to room, temperature slides were washed 3x with PBS and liquid blocked with a marker. 200ul of peroxidase (covering the tissue) was added and incubated for 10 minutes before being washed off with PBS. Protein block background (with background sniper from Biocore Medical) was added and incubated for 15 minutes at room temperature. Slides were rinsed in PBS for 1-2 mins. Slides were then blocked for 60 minutes with goat serum, and the primary antibody was then added according to manufactures recommendations and incubated overnight. After overnight incubation, slides were washed with TBST/Tween solution three times for 1 minute. The secondary antibody was added and incubated at room temperature for 35 mins after which it was washed off 3x with PBS. DAB chromogen (diluted 1:2 in DAB buffer) was added and incubated for 3 minutes. Hematoxylin counterstain was then applied for 1 minute and removed. Slides were then dehydrated using graded alcohol and xylazine (95% ETOH 3mins, 100% ETOH 2x 3 mins, Xylene 3x 5mins), and a coverslip was applied using xylene based mounting medium.

### ChIP Sequencing/Bioinformatics

Cells exposed to either DMSO or TMZ for the desired amount of time were washed twice with sterile PBS and then exposed to 1% PFA (Sigma) for crosslinking. After crosslinking for 15 minutes, the reaction was stopped with 2.5 M glycine (Sigma), and cells were rocked for an additional 5 minutes. Cells were then collected by scraping using minimal amounts of PBS to wash cells off the dish. The scrapings were collected, and pellets of cells were flash-frozen and sent to Zymogen research corporation to undergo ChIP-sequencing per their established protocols. Sequencing data was returned to us and processed bioinformatically after FastQC determined sequencing to be satisfactory. Alignment to the reference genome Hg38 was done using Tophat2, and peak calling were done using MACS2 software with a p-value threshold set to 0.05. Peaks were initially visualized using IGV, and graphical visualizations were made by using a combination of both ChIPSeeker2 and pygenometracks^25^.

### Metabolite isolation and Liquid Chromatography-Mass Spectrometry (LCMS) Profiling

To determine the relative abundances of intracellular metabolites, extracts were prepared and analyzed by LC-MS/MS. Briefly, for targeted steady-state samples, metabolites were extracted on dry ice with 4-mL 80% methanol (−80°C), as described previously^26^. Insoluble material was pelleted by centrifugation at 3000 g for 5 min, followed by two subsequent extractions of the insoluble pellet with 0.5-ml 80% methanol, with centrifugation at 20,000g for 5 min. The 5-ml metabolite extract from the pooled supernatants was dried down under nitrogen gas using the N-EVAP (Organomation, Inc, Associates). 50% acetonitrile was added to the dried metabolite pellets for reconstitution. Samples solution was then centrifuged for 15 min at 20,000 g, 4°C. The supernatant was collected for LC-MS analysis. Hydrophilic Metabolites Profiling: Samples were analyzed by High-Performance Liquid Chromatography and High-Resolution Mass Spectrometry and Tandem Mass Spectrometry (HPLC-MS/MS). Specifically, the system consisted of a Thermo Q-Exactive in line with an electrospray source and an Ultimate 3000 (Thermo) series HPLC consisting of a binary pump, degasser, and auto-sampler outfitted with an Xbridge Amide column (Waters; dimensions of 4.6 mm × 100 mm and a 3.5 µm particle size). The mobile phase A contained 95% (vol/vol) water, 5% (vol/vol) acetonitrile, 20 mM ammonium hydroxide, 20 mM ammonium acetate, pH = 9.0; B was 100% Acetonitrile. The gradient was as following: 0 min, 15% A; 2.5 min, 30% A; 7 min, 43% A; 16 min, 62% A; 16.1-18 min, 75% A; 18-25 min, 15% A with a flow rate of 400 μL/min. The capillary of the ESI source was set to 275°C, with sheath gas at 45 arbitrary units, auxiliary gas at five arbitrary units, and the spray voltage at 4.0 kV. In positive/negative polarity switching mode, an m/z scan range from 70 to 850 was chosen, and MS1 data were collected at a resolution of 70,000. The automatic gain control (AGC) target was set at one × 106, and the maximum injection time was 200 ms. The top 5 precursor ions were subsequently fragmented, in a data-dependent manner, using the higher energy collisional dissociation (HCD) cell set to 30% normalized collision energy in MS2 at a resolution power of 17,500. Data acquisition and analysis were carried out by Xcalibur 4.1 software and Tracefinder 4.1 software, respectively (both from Thermo Fisher Scientific).

### In Vivo Metabolite Tracing

Using BLI, it was determined that mice had a well-established tumor but were still healthy enough to undergo longer-term isoflurane exposure. Mice were anesthetized using Isoflurane and placed into a tail vein exposure rig (Braintree Scientific). Using butterfly 27 gauge needles, the vein was accessed, and the isotope of choice was administered to the animal. The isotope solution was dissolved into sterile saline, and the infusion was given as an initial bolus of ∼100ul over 90 seconds, and a final infusion of ∼250ul infused at 3ul/min rate for 2 hours using an infusion pump (Braintree Scientific). Tails of the mice were observed during the entire procedure to ensure needle placement within the vein was maintained. Once the infusion was complete, the mice were immediately sacrificed, desired tissue was then excised and flash frozen until metabolite extraction could take place. For metabolite extraction, samples were homogenized and suspended in 80% MeOH (sigma) cooled to −80C. After subsequent spins and resuspensions of sample pellets in ice-cold MEOH, the cell pellets were saved for later protein quantification. At the same time, the metabolite containing supernatant was subjected to LN2 speed vac drying. Pellets resulting from the drying were stored at −80 until they were sent to a proteomics core for analysis using LC-MS.

### Extreme limiting dilution assay

Following trypsinization, cells are diluted to a concentration of 1 cell/uL in neurobasal media supplemented with glutamine, N2 supplement, B27 supplement, EGF, FGF, and 1% Antibiotic Antimycotic mixture (all components of neurobasal media provided by Gibco). Cells are plated in an untreated 96-well tissue culture plate (VWR) at eight different densities (3 cells/well to 200 cells/well) in replicates of 12. Neurosphere formation is assessed seven days after plating. The analysis is done using the R package provided by the Walter+ and Eliza Hall Institute of Medical Research.

### U-^14^C-glycine, ^3^H-hypoxanthine, ^14^C-guanosine and ^3^H-guanine incorporation into RNA and DNA

Cells (∼80% confluent) were incubated for 15 hours, then treated and labeled as indicated in the figures. Cells were labeled with 2-μCi of either U-14C-glycine, 3H-hypoxanthine, and 3H-guanine. Cells were harvested, and RNA or DNA was isolated using Allprep DNA/RNA kits according to the manufacturer’s instructions and quantified using a spectrophotometer. 70 μl of eluted DNA or 30 μl of eluted RNA were added to scintillation vials, and radioactivity was measured by liquid scintillation counting and normalized to the total DNA or RNA concentrations, respectively. All conditions were analyzed with biological triplicates and representative of at least two independent experiments.

### Statistics

All statistics were performed by accompanying analysis software when indicated or by graphpad prism software version 8.0 if not indicated. Survival statistics were compared between multiple groups using Bonferroni correction applied to Log-Rank tests. Generally, T-Tests and ANOVAs (one and two way) were used to preform analyses and all p values reported are adjusted for multiple comparisons when appropriate.

## Results

### EZH2 expression is increased in the BTIC compartment during chemotherapy

Several reports have demonstrated the significance of PRC2/EZH2 activity in gliomagenesis as well as maintenance of the BTIC niche ^16, 19^. To investigate its role in promoting chemotherapy-induced plasticity and chemoresistance in GBM, we examined the expression status of the catalytic subunit of the PRC2 complex, EZH2, in both PDX lines as well as GBM cell lines. Fluorescence-activated cell sorting (FACS) analysis revealed that a clinically relevant dose ^27–29^ of TMZ increased EZH2 levels while increasing the BTIC frequency as defined by CD133 (Sup. Fig. 1A & B). Moreover, GBM cells co-express EZH2 and CD133 in a time-dependent manner post-TMZ exposure. Immunoblot analysis of tumor tissues isolated from two different subtypes of PDX tumors confirms elevated EZH2 expression during therapy (Fig. 1B). FACS analysis demonstrated a significant increase in EZH2 positive (+) cells in two different models following therapy (Fig. 1 C & D & E; One Way ANOVA adjusted p-values, *=p<.05, ***=p<.001). Although EZH2 expression was up generally, we saw the most considerable increase in CD133+ BTIC compartment both during and post TMZ therapy. An shRNA mediated EZH2 knockdown PDX line (Sup. Fig. 1C) shows that in the absence of EZH2, TMZ-mediated increase of the CD133+ BTICs was significantly inhibited (Fig. 1F; One Way ANOVA adjusted p-values sh Control (shC) vs. sh3 p<.05, shC vs. sh4 p<.001). To validate this finding in vivo, we used 3-Deazaneplanocin (DZNep), an adenosylhomocysteine inhibitor previously reported to inhibit EZH2 ^16^. In the subcutaneous PDX model, DZNep significantly reduce post-therapy CD133+ cells, the double-positive CD15+CD133+ cells (Fig. 1G; two-way ANOVA adjusted p values CD133+ DMSO vs. TMZ p<.001, CD133+CD15+ DMSO vs. TMZ p<.001), as well as CD133+ cells expressing BTIC-specific transcription factor, SOX2 (Fig. 1G; two-way ANOVA adjusted p-values CD133+SOX2+ TMZ vs. EZH2 Inhibitor p<.001, TMZ vs. TMZ+EZH2 Inhibitor p<.001). Previous reporting demonstrated that Hypoxia-inducible factors (HIFs) are critical for maintenance of the BTIC niche and that HIF-1α (HIF1A) can directly modulate EZH2 expression via promoter binding ^30, 31^. We have shown that TMZ therapy increases the BTIC frequency in a HIF dependent manner ^20, 32^. Now we show that the TMZ-mediated increase in HIF1A is specific to the CD133+ BTIC compartment (Sup. Fig. 1D; One Way Anova adjusted p-values ***=p<.001). To examine the regulatory effects on EZH2, shRNA-mediated knockdown of HIFs were generated (Sup. Fig. 2C; One Way ANOVA adjusted p-values **=p<.01, ***=p<.001). Knocking down HIF1A, but not HIF2A, results in a significant reduction in the TMZ-mediated BTIC expansion (Fig. 1H; One Way ANOVA adjusted p values shControl_HIF1 sh1 p<.001, shControl_HIF2 sh3 p is ns). Moreover, HIF1A expression was positively correlated with EZH2 expression in patient GBM samples (Sup. Fig. 2A, GlioVis^33^ CGGC Dataset Pearsons correlation r=0.42, Rembrandt r = 0.52) Regional gene expression patterns of HIF1 and EZH2 were also similar with expression lowest in the leading edge of the tumor and significantly higher in the perinecrotic zone and pseudopalisading cells when compared. (Suppl. Fig 2B, Tukey’s pairwise comparisons). Finally, immunofluorescent (IF) analysis of mouse brain with PDX revealed colocalization of HIF1A and EZH2 expression (Fig. 1I).

**Figure 1:**
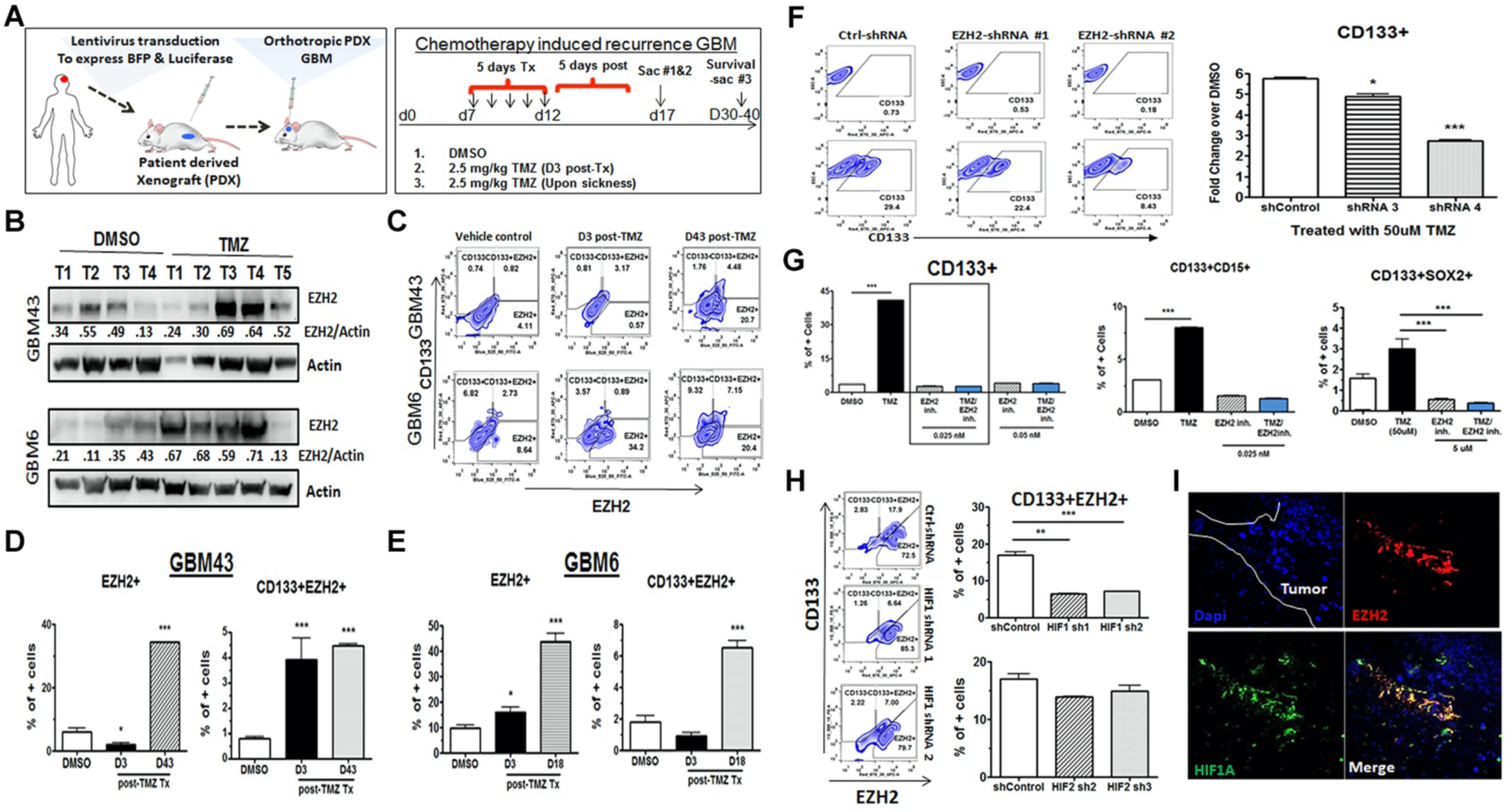
EZH2 expression is increased in the BTIC compartment during TMZ-based chemotherapy. **A**. Schematic showing establishment of a blue fluorescent protein (BFP) and luciferase expressing PDX line as well as the treatment regimen for establishment of recurrent tumors. **B**. Western blot analysis on tumor tissue isolated from intracranial PDX tumor treated with DMSO or TMZ. Numbers are relative pixel intensity of EZH2 bands normalized to actin (n=4-5). **C**. Intracranial PDX tumor analyzed by Fluorescence-Activated Cell Sorting (FACS) with HLA staining for human cells, BTIC marker CD133, and EZH2 during therapy (day 3/D3) and in recurrence (D43/post therapy when mice show signs of disease). **D**. Bar graphs displaying percent of cells staining for either EZH2 alone or EZH2+CD133+ double positive cells at days 3, 18, and 43 post therapy for GBM43 (n=3); **E**. Same as before for GBM6 (n=3). **F**. Representative FACS plots from GBM6 transduced with lenti-vectors carrying scramble control (shControl) and two different shRNA for EZH2. Right, bar graph representing percent of cells residing in CD133+ gate. **G**. Bar graph representing FACS data displaying percent of cells showing CD133+, CD133 & CD15+, or CD133 & SOX2+ double positive, GBM43 PDX cells treated with DMSO, TMZ, EZH2 inhibitor, or combination *in vivo* flank model as EZH2 inhibitor did not cross the blood brain barrier. **H**. FACS analysis plots and bar graph displaying CD133 and EZH2 staining across shControl and *HIF1A* or *HIF2A* knockdown with two shRNAs. Numbers represent percent of cells residing within specific gate. Bar graphs represent FACS data showing percent of cells expressing both CD133 & EZH2 across the shRNA treatment with either HIF1A or HIF2A knockdown using multiple shRNA **I**. Representative images from orthotopic PDX tumor stained for HIF1A and EZH2. All error bars in graphs depict 3 technical FACS replicates for each animal, each analysis is validated in at least 3 different animals and represent mean + SD. *p<.05 ***p<.001.

### Ciliary protein ARL13B is a downstream target of EZH2 and its expression negatively correlated with GBM patient’s prognosis

We postulate the PRC2/EZH2 complex alters the expression of genes necessary for TMZ-induced cellular plasticity by modulating the chromatin landscape, ultimately driving therapeutic resistance. To identify potential targets of EZH2, we employed gene expression analysis with two subtypes of PDX treated with TMZ and with or without EZH2 inhibitor. The most highly enriched target was ARL13B, as its expression was altered over 6-fold in the presence of DZNep as opposed to TMZ treatment alone (Fig. 2A; p<0.05). ARL13B is an ADP-ribosylation factor family small GTPase primarily described as a ciliary protein; however, its role in gliomagenesis is understudied. To investigate this, we first examined the relationship between EZH2 and ARL13B in the context of GBM. Gliovis data show a positive correlation between ARL13B and EZH2 expression in GBM, astrocytoma and oligodendroglioma (Fig. 2E; R=0.5, p<0.005; R=0.32, p<0.005; R=0.53, p<0.005), but not in non-tumor tissue (Fig. 2E; R=0.-27, p<0.16). Given EZH2’s global regulatory role, we did a genome-wide Chromatin immunoprecipitation sequencing (ChIP-seq) analysis for EZH2 and H3K27me3 marks during therapy (Sup. Fig. 3A). These data indicated a possible interaction with the Stat3 pathway, an established oncogenic driver of gliomagenesis (Sup. Fig. 3B, C & D)^34^. As the EZH2-Stat3 axis has been previously reported to be important contributors to gliomagenesis^19, 35^, such observation serves as validation for our screen. The screen also indicated EZH2 could directly bind to one of the established ARL13B enhancer regions (Ch3:93470260-93470889)^36^ and this interaction was significantly decreased four days post-TMZ therapy as compared to control (Fig. 2B; 2.5 fold decrease in expression day 1 (D1): p<1E-47, day 4 D4: p<1E-17)). The presence of H3K27me3, EZH2/PRC2’s canonical silencing mark, correlated both spatially and temporally with the binding of EZH2 within the ARL13B enhancer.

**Figure 2:**
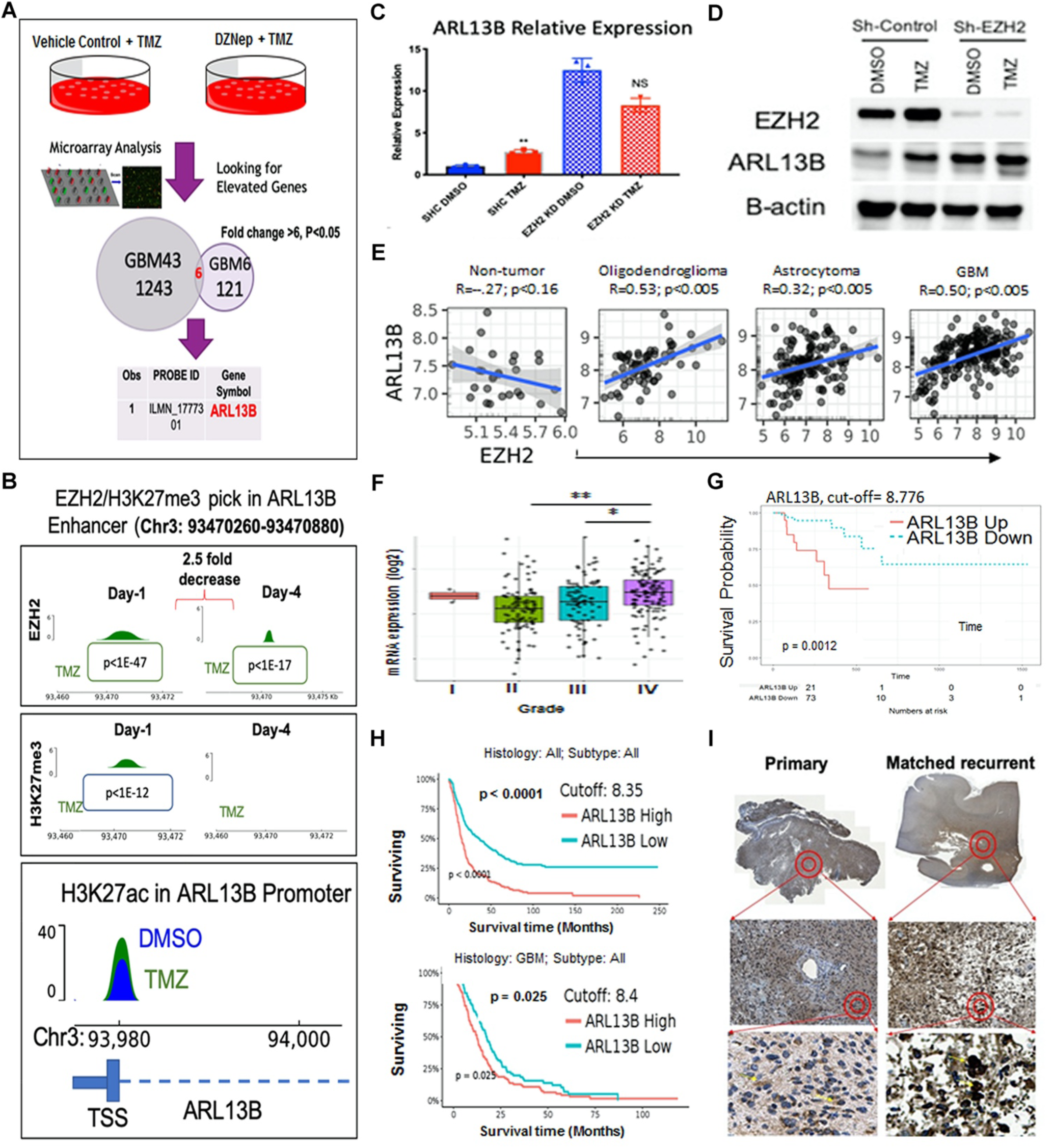
Ciliary protein *ARL13B* is a downstream target of *EZH2* and its expression is negatively correlated with GBM patient prognosis. **A**. Schematic diagram of experimental design where GBM43 and GBM6 were treated with either vehicle control or DZNep (0.05 nM) in combination with TMZ (50μM) for 48 hours. Gene expression analysis between the different subtypes of GBM identified *ARL13B* as one of the top genes for which expression was altered in the absence of EZH2 activity (fold change >6, p<0.05). **B**. Genome-wide Chromatin Immunoprecipitation Sequencing (ChIP-Seq) analysis is performed on GBM43 cells at Day 1 and Day 4 after TMZ (50μM) for enrichment of EZH2, H3K27me3 and H3K27ac marks. P-values from Macs2 software called over input control. **C.** Quantitative polymerase chain reaction (qPCR) analysis demonstrating that *ARL13B* expression increases after 4 days of TMZ treatment as compared to vehicle control. Error bars represent mean + SD of 3 technical qPCR replicates. Each experiments repeated minimum twice. **D**. Immunoblot analysis corroborating qPCR data from C at the protein level at 4 days post TMZ treatment. **E**. TCGA data demonstrating significant positive correlation between *EZH2* expression and *ARL13B* expression in oligodendroglioma, astrocytoma, and GBM but not in normal brain. **F.** TCGA data showing mRNA expression of *ARL13B* increasing significantly with tumor grade, **G**. probability of survival decreasing with higher expression of *ARL13B*, and **H.** time to median survival significantly decreasing with high expression of *ARl13B* in all brain tumor (top) and only in GBM (bottom). Data were gathered using optimal cutoff. **I**. GBM patient tissue samples stained for *ARL13B* in matched pairs of primary and recurrent tumors demonstrating increased nuclear staining for *ARL13B* upon recurrence (n=10). * p<.05 ** p<.01.

These binding dynamics correlated with an increase in H3K27ac at the transcription start site of ARL13B four days post TMZ therapy (Fig. 2B; bottom panel, Sup. Fig. 4A; table 2.1-fold enrichment TMZ/DMSO p<2.1E-22, q<2.6E-18), which resulted in increased ARL13B at both the transcript (Fig. 2C, one-way ANOVA adjusted p=.0069, n=3 replicates) and protein levels during TMZ therapy(Fig. 2D; Sup. Fig. 4B). Similarly, the shRNA knockdown of EZH2 also elevated the levels of ARL13B mRNA and protein (Fig. 2C, D & Sup. Fig. 4B, One-way ANOVA adjusted p =.0121 (DMSO), adjusted p=.0288 (TMZ)). These data indicated that EZH2 could directly bind to the ARL13B regulatory enhancer region, and such binding is decreased during TMZ therapy, allowing GBM cells to induce ARL13B expression in a controlled manner.

We also examined GBM patients’ data using Gliovis visualization tools to investigate the role of ARL13B in gliomagenesis. Data indicated that ARL13B expression increased concurrently with tumor grade (Fig. 2F; TCGA Rembrandt, ANOVA test Grade II vs. IV adjusted p<.0001. Grade III vs. IV adjusted p<.0001), and its elevated expression is negatively correlated with overall survival (Fig. 2G, right, Log-Rank cutoff = 8.776 p=.0012). Importantly, high ARL13B expression was associated with accelerated relapse and recurrence (Fig. 2G & H Log-Rank cutoff 8.35 p<.0001, cutoff 8.4 p<.025). We profiled ten matched primary and recurrent GBM’s from Northwestern Nervous System Tumor Bank are compared for ARL13B expression via immunohistochemistry (IHC). This analysis by our board-certified neuropathologist revealed that ARL13B is well expressed in both the primary and recurrent tumors but that a nuclear localization shift seemed to occur in the matched recurrent slides. (Fig. 2I, Sup. Fig. 4C).

### ARL13B regulates cellular plasticity and contributes to tumor engraftment in vivo

As data indicated that during TMZ therapy, EZH2 promotes plasticity as well as regulates ARL13B expression, we next investigated if ARL13B itself can be involved in promoting plasticity in GBM. Knocking down ARL13B significantly reduced the expression of several stemness transcription factors such as SOX2 and Oct4 (Fig. 3A). Knockdown also reduced GBM cell stemness measured by extreme limiting dilution assay (ELDA) (Fig. 3B, Left, Left, BTIC frequency for Control 1/28, ARL13B KD 1/65.1, p<0.002). The loss of stemness can be rescued by re-expressing ARL13B in KD cells (Fig. 3B, Right, BTIC frequency for KD 1/72.8, Rescue 1/53.6 p>0.05). To examine if the cell fate state influenced the expression of ARL13B, we cultured PDX lines in stem cell media (NEB, neural basal media with 10 ng/ml EGF and FGF) or differentiation condition media containing 1% fetal bovine serum^32, 37^. Immunoblot analysis indicated that when PDX lines cultured in NEB, the expression of ARL13B was significantly elevated in three different subtypes of GBM (Fig. 3C and Sup. Fig. 5).

**Figure 3:**
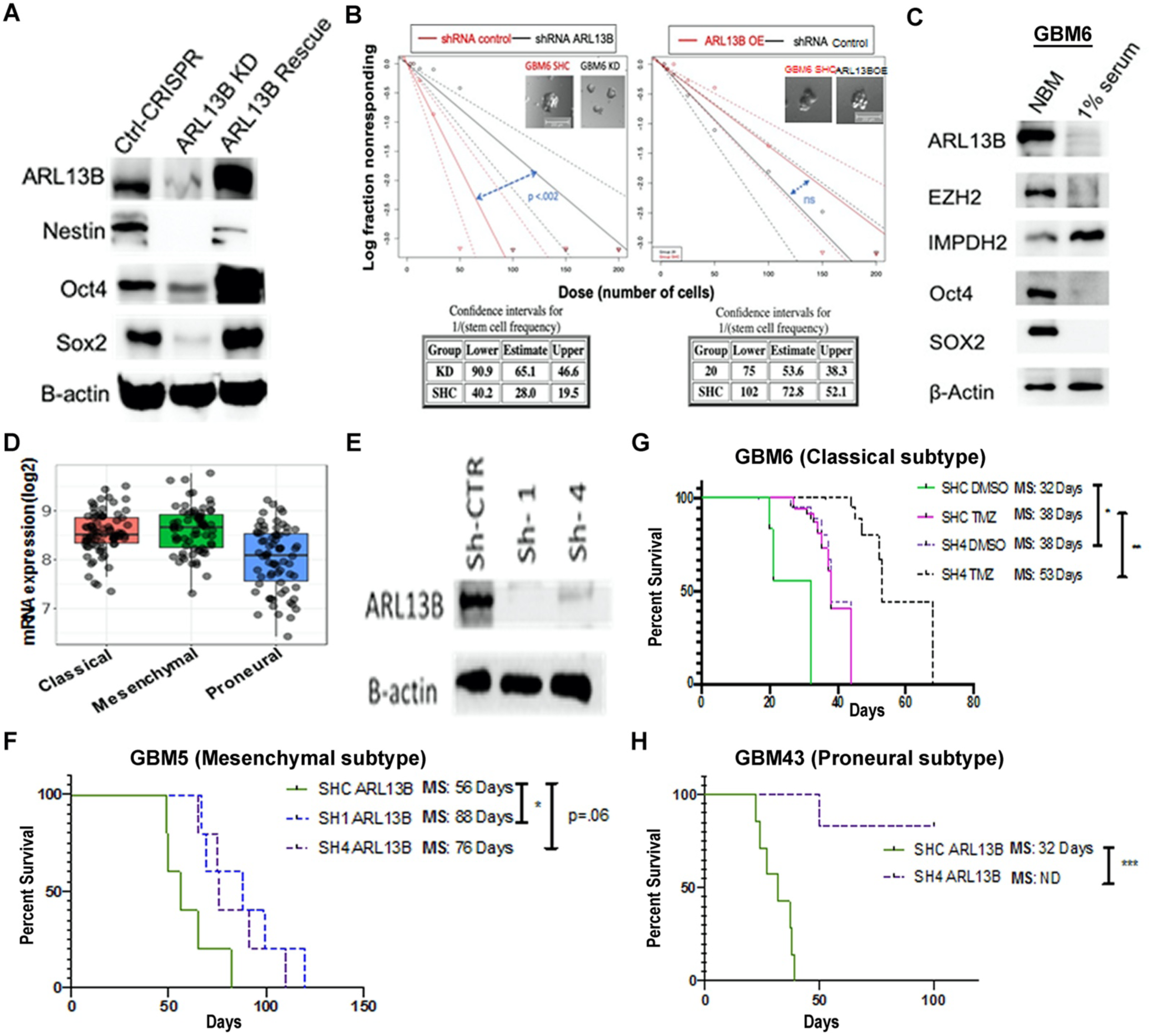
*ARL13B* regulates cellular plasticity and is required for tumor engraftment in patient derived xenograft models. **A**. Western blot analysis of canonical stem cell markers in U251 GBM cells with wild type expression, CRISPR-mediated deletion, and lentiviral-mediated rescue of *ARL13B*. **B**. Extreme limiting dilution analysis assay on cells with CRISPR KD of ARL13B or lentiviral rescue of ARL13B compared to nontargeted guide RNA. Error bars demonstrate 95% CI of stem cell frequency p-value determined from 12-well replicates of plate counting in provided software. **C.** Immunoblot analysis showing differential expression of ARL13B, EZH2 an IMPDH2 as well as canonical stem cell markers SOX2 and Oct4 in the neurobasal (10 ng/ml EGF, 10 ng/ml FGF) or differentiation media (1% FBS). **D**. TCGA data demonstrating mRNA expression of *ARL13B* across GBM subtypes. **E**. Immunoblot analysis showing shRNA mediated knockdown efficiency of *ARL13B* in GBM5 PDX line. **F.** *In vivo* engraftment analysis after removal of *ARL13B* using shRNA from mesenchymal (**F**, GBM5), classical (**G**, GBM6) and proneural (**H**, GBM43) subtypes of PDX GBM (n=6 mice per group, 50% male and female mice). * p<.05, ** p<.01, *** p<.005.

In GBM patient samples, ARL13B mRNA expression was very similar in the classical and mesenchymal subtype. However, the proneural subtype expression was significantly lower (Fig. 3D; Gliovis; One-Way ANOVA adjusted p<.0001). To investigate the role of ARL13B in GBM growth *in vivo*, the shRNA approach was employed to knock down its expression in all three subtypes of PDX GBM (Fig. 3E). Knocking down of ARL13B resulted in a 57% improvement of median survival (MS) in the mesenchymal subtype (Fig. 3F; Log-Rank adjusted for multiple comparisons shC vs. sh1 p=.02, shC vs. sh4 p=0.06); a 72% improvement of MS in classical subtype GBM6 (Fig. 3G; Log-Rank p-adjusted=.0002); and completely blocked tumor formation in proneural GBM43 subtype (Fig. 3H; Log-Rank p-adjusted=.0004).

### Removal of ARL13B sensitized GBM cells to TMZ

Our initial experiments identified ARL13B as a downstream target of EZH2 during TMZ therapy, therefore, to investigate if removal of ARL13B sensitized GBM to TMZ therapy *in vivo*, we established PDX lines with shRNA mediated ARL13B knockdown and intracranially injected these into nude mice. Following tumor engraftment (7 days post-implantation), animals were treated with a 2.5mg/kg dose of TMZ^38^. Removal of ARL13B significantly sensitized mesenchymal PDX GBM5 with approximately 25% increase in MS (Fig. 4A, shC+DMSO vs. sh4+DMSO adjusted Log-Rank p=.07; shC+TMZ vs. sh4+TMZ adjusted Log-Rank p=.05; sh4+DMSO vs. sh4+TMZ adjusted Log-Rank p=.102), aa well as classical PDX GBM6, approximately 21% increase in MS, demonstrating a role for ARL13B in response to TMZ therapy (Fig. 4B, shC+DMSO vs. sh4+DMSO adjusted Log-Rank p=.025; shC+TMZ vs. sh4+TMZ adjusted Log-Rank p=.004; sh4+DMSO vs. sh4+TMZ adjusted Log-Rank p=.0021). Knocking down nearly abolished the engraftment capacity of the proneural subtype GBM43 (Fig. 4C, shC+DMSO vs. sh4+DMSO adjusted Log-Rank p=.0004, shC+TMZ vs. sh4+TMZ adjusted Log-Rank p=.0003), preventing an assessment of the effect of TMZ. However, KD of ARL13B.

**Figure 4:**
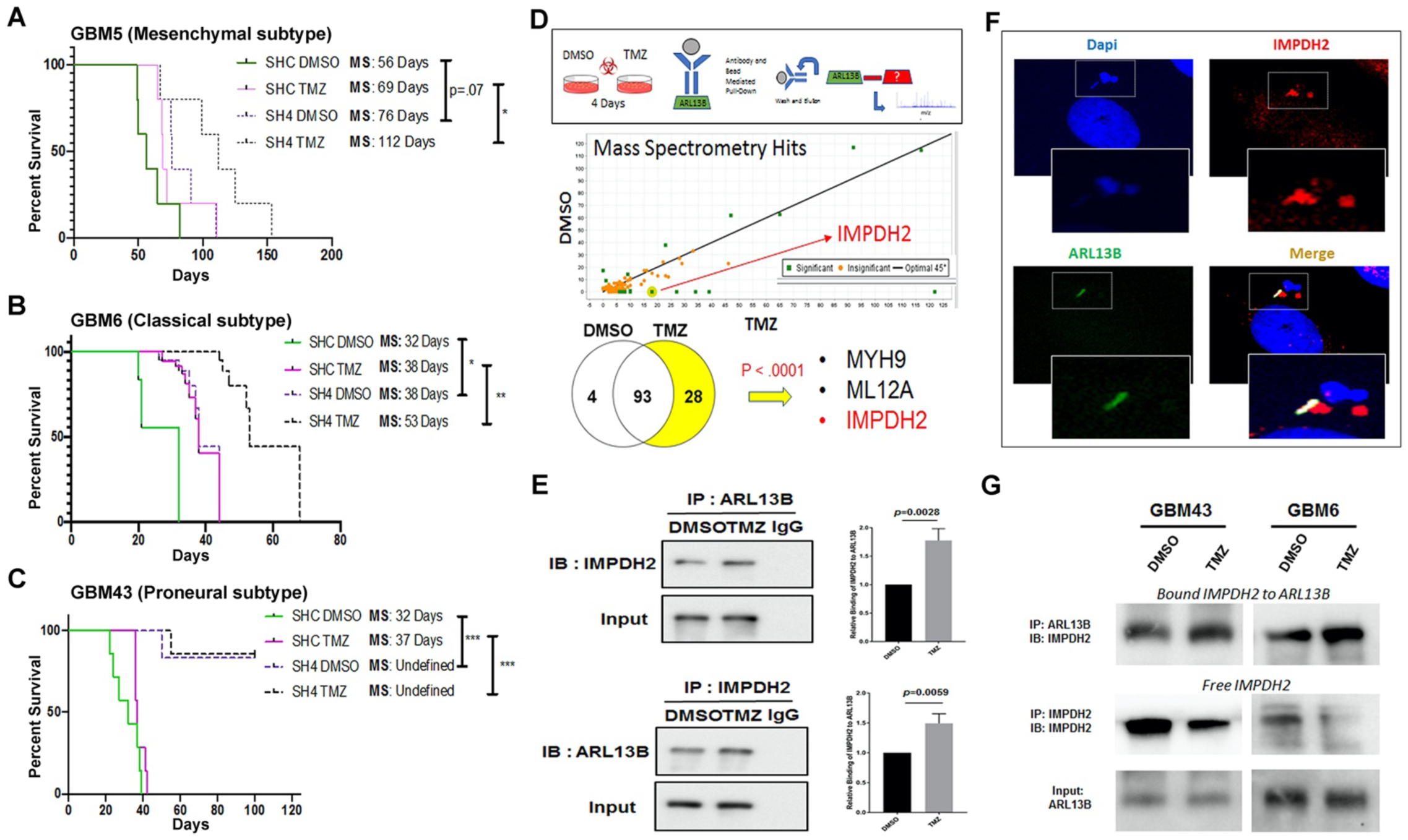
Loss of *ARL13B* increases *in vivo* sensitivity to TMZ and *ARL13B* interacts with *IMPDH2* during TMZ therapy. **A.** Kaplan-Meier curves for endpoint survival analysis evaluating the role of ARL13B in TMZ sensitivity. shRNA-mediated knockdown of PDX lines **A.** GBM5, **B.** GBM6, and **C.** GBM43 were engrafted intracranially and treated with TMZ (2.5 mg/kg) for 5 consecutive days beginning 7 days after tumor implantation (n=6) and mice are evaluated for endpoint survival. **D**. Schematic representing experimental flow of antibody pulldown mass spectroscopy analysis. p-values determined by Scaffold proteomics software. **E.** Immunoprecipitation(IP) using anti-IMPDH2 or anti-ARL13B antibodies with an IgG control during DMSO or TMZ therapy to evaluate ARL13B IMPDH2 interaction. Error bars represent densitometry quantification from biological triplicate IPs. **F**. Immunofluorescent co-staining with antibodies against IMPDH2 and ARL13B in GBM43 line. **G**. Top, schematic representation of the different protein domains of ARL13B tagged with Flag and expressed in HEK293 cells in order to determine the binding domain of the IMPDH2 interaction. IP was done using anti-HA epitope antibody followed by immunoblot with corresponding antibody. **H**. Representative IP experiment demonstrating the levels of free and bound *IMPDH2* protein during TMZ therapy across GBM6 and 43. Samples were serially IPs’ as described in methods to assay free and bound forms of the protein.

### IMPDH2 interacts with ARL13B during TMZ therapy in PDX GBM model

ARL13B is known to localize around primary cilia, a sensory organelle found on eukaryotic cells known to function as an hub for cellular signaling, including sonic hedgehog signaling (Shh)^39, 40^. Shh signaling is known to contribute to various human malignancies, including the childhood brain tumor medulloblastoma^41–43^. In PDX GBM, the length of ARL13B positive cilia, as well as the number of ciliated GBM cells, increased significantly after TMZ exposure (Sup. Fig. 6A, 7A & 7B). However, total Shh signaling, as well as the suppressor of Shh signaling Sufu, remained unchanged (Sup. Fig. 6B & C). Furthermore, the elimination of cilia pharmacologically did not yield a decrease in cancer cell proliferation in combination with TMZ (Sup. Fig 7C). This data indicated to us a possible alternative mechanism of ARL13B-mediated TMZ resistance in GBM independent of Shh.

To examine other potential mechanisms, we performed immunoprecipitation (IP) with the ARL13B antibody, followed by liquid chromatography-mass spectrometry (LC-MS). This analysis identified inosine monophosphate dehydrogenase 2 (IMPDH2), a key rate-limiting enzyme for purine biosynthesis^44, 45^, as a novel interacting partner of ARL13B (Fig. 4D; p<.0001 calculated by Scaffold Viewer software n=2 replicates). Moreover, this interaction is significantly augmented during TMZ therapy (Fig. 4E; n=3 biological replicates ARL13B IP: p=.0028, IMPDH2 IP: p=.0059). The interaction was further validated in different PDX lines (Sup. Fig. 8C) as well as in breast cancer cells (Sup. Fig. 8D). Time course analysis of post-therapy expression indicated subtype-specific increases is IMPDH2, EZH2, and ARL13B during therapy with pronounced increases at day 8 in GBM 43 and 6 (Sup. Fig. 8A & B). Immunofluorescent (IF) analysis in PDX lines revealed the co-localization of both proteins in the cytoplasm and around primary cilia (Fig. 4F). Finally, sequential IP was performed to examine the dynamics of this interaction and observed that within 24 h after TMZ exposure the free IMPDH2 level was reduced as the ARL13B-IMPDH2 interaction is increased (Fig. 4G).

### ARL13B and IMPDH2 interaction regulates purine synthesis through the consumption of hypoxanthine

Purines can be built in the cell using one of two pathways: first, the energy-demanding de-novo pathway where the purine is generated within the cell from a ribose ring^46^. Second is a more energy-efficient salvage pathway where purines can be recycled from the environment^47, 48^. All cells can perform both types of synthesis; however, the brain typically utilizes the energy-efficient salvage pathway over the energetically taxing de novo pathway^49^. IMPDH2 is the rate-limiting enzyme for IMP-XMP-GMP conversion, essential for both de novo as well as salvage pathway. As the IMPDH2-ARL13B interaction has never been reported in the literature, we hypothesized that this binding event might influence GBM cells’ ability to synthesize purines. Our initial ChIP-seq data revealed purine biosynthetic enzymes significant changes in both relative expressions, as well as the transcription activation marks H3K27ac at the corresponding TSS after 4 days of TMZ therapy (Sup. Fig. 9). To specifically investigate the role of the IMPDH2-ARL13B interaction, we employed radiolabeling tracing into nucleic acids using [^14^C]-glycine to measure the de novo purine pathway, [^3^H]-hypoxanthine that provides a direct measurement of de novo GMP synthesis through IMPDH, and [^14^C]-guanosine which is directly converted to GMP independently of IMPDH activity via the purine salvage enzyme hypoxanthine guanine phosphoribosyltransferase (HGPRT). Measurement of radiolabeled isotope incorporation within DNA and RNA indicated that TMZ decreased the amount of salvage activity ([3H]-hypoxanthine incorporation) while having no significant effect on de novo synthesis ([^14^C]-glycine) in DNA (Fig. 5A; 2-way ANOVA Tukey adjusted p=.003). However, in cells lacking ARL13B, there was a 4-6 fold increase in salvage activity ([^3^H]-hypoxanthine incorporation), which was unaffected by TMZ treatment. Knockdown of ARL13B also significantly reduced the amount of de novo synthesis ([^14^C]-glycine incorporation; Fig. 5A & B; DNA Salvage Tukey adj p,.0001, RNA Salvage Tukey adj p<.0001, DNA de-novo Tukey adj p=.002). Conversely, IMPDH2-independent salvage ([^14^C]-guanosine incorporation) was increased in response to ARL13B knockdown, although to a much lesser extent than IMPDH2 dependent salvage ([^3^H]-hypoxanthine) (Fig. 5A & B, DNA Tukey adj p<.0001, RNA Tukey adj p=.6). These results demonstrate how the knockdown of ARL13B impairs GBM cells ability to utilize de-novo synthesis, instead forcing salvage pathway utilization. These observations were next validated in PDX lines GBM 43 & 6 using shRNA-based knockdown of ARL13B (Fig. 5C & D; One-way Adjusted ANOVA 43 DNA p=.0003, p=.0008, 43 RNA p,>0001 p=.006 6 DNA p=.004).

**Figure 5:**
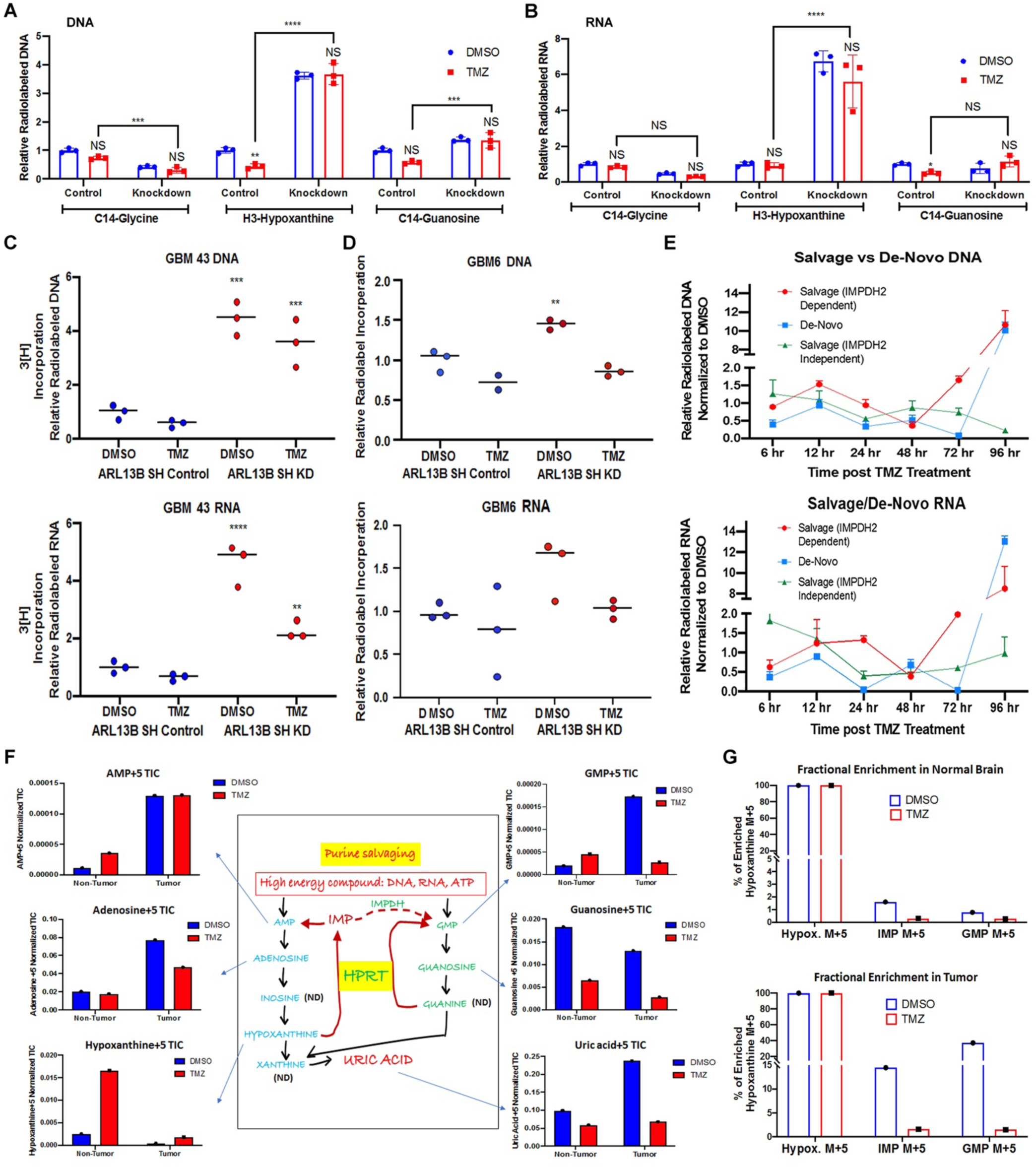
ARL13B and IMPDH2 interaction functions as a negative regulator of purine salvaging and TMZ therapy alters purine biosynthesis pathways. **A**. Radiolabeled tracing of the three major biosynthetic pathways measured by the corresponding radiolabeled metabolite incorporation at the DNA level with or without ARL13B, in the absence and presence of TMZ (50μM). Graph depicts relative incorporation of specific radiolabel normalized to control DMSO condition. **B**. Same experiments as described before but measuring incorporation in RNA. **C**. Radiolabeled tracing of GBM43. Graph depicts relative ^3^H incorporation in the DNA across cells with and without ARL13B during TMZ therapy. **C.** Same experiment as described before but using GBM6 PDX line. **E**. Radiolabeled tracing of different purine biosynthesis pathways after 6, 12, 24, 48, 72 and 96 hours post TMZ exposure in GBM43. Data presented as normalized to DMSO incorporation from matched timepoint. **F.** Bar graph representing IMPDH2 activity assay data measured in nM/min/ug. Assay done on U251 cells expressing endogenous ARL13B (WT), CRISPR-mediated knockdown of ARL13B (KD), ARL13B rescue from knockdown, and ARL13B overexpression (OE). **G**. IMPDH2 activity (as measured above) in U251 cells with knockdown of *ARL13B*, rescued with either wild-type ARL13B (WT) or a mutated GTP binding domain (GTP null). **H**. Mice bearing GBM43 PDX tumors were infused with labeled hypoxanthine (13C5) for two hours through systemic tail vein injection. Tissues were collected from intracranial tumor, contralateral normal brain, and subject to analysis by LC-MS. Bar graphs represent *in vivo* tracing of metabolites from mice bearing a GBM43 PDX tumor. All counts are represented as normalized total ion counts (TIC) and were isolated from tumor or non-tumor tissue +/- TMZ. **I** Fractional enrichment for Hypoxanthine, IMP, & GMP represented as a bar graphs measured from samples of normal brain tissue. **J**. Same experiment as above but representing fractional enrichment in tumor tissue. Enrichment was calculated from protein-normalized TIC and normalized to 13C hypoxanthine incorporation. All error bars in experiments represent technical replicates and display mean + SD. ** p<.01 ***p<.001 **** p<.0001.

We next examined how purine biosynthesis was altered in response to a physiological dose of TMZ over time. We observed a higher utilization of salvage synthesis ([^3^H]-hypoxanthine incorporation) carrying up to 48 hours where de novo synthesis ([^14^C]-glycine incorporation) briefly outpaced it^27, 28, 50^. After 48 hours, both IMPDH2-dependent de novo synthesis and IMPDH2-dependent salvage synthesis rebounded and even drastically increased roughly 10-fold by 96 hours. However, IMPDH2-independent salvage synthesis ([^14^C]-guanosine incorporation) remained relatively low (Fig. 5E).

### TMZ downregulates purine salvage in vivo

Our data strongly suggested that TMZ is altering the purine synthesis pathways used in glioma. To analyze *in vivo* effects, we utilized heavy isotope-labeled hypoxanthine (^13^C5) to quantify the purine biosynthesis flux through the IMPDH2-dependent salvage pathway. Mice bearing GBM43 PDX tumors are infused with labeled hypoxanthine for two hours through systemic tail vein infusion. LCMS examination was conducted post-infusion for isotope incorporation in tumor tissue, contralateral normal brain, and liver tissue (Sup Fig. 10 A & B). Results demonstrated that the tumor region contained a higher level of labeled AMP, adenosine, and Uric acid as compared to cells from the non-tumor area (Fig. 5F). Furthermore, mice harboring subcutaneous GBM 43 demonstrated increased fractional enrichment when compared back to normal brain and tumor brain within the same mouse, indicating that microenvironment plays a role in modulating purine biosynthesis (Sup. Fig. 10C).

Interestingly, the levels of labeled hypoxanthine, the first metabolite injected, were much lower within the tumor when compared to the non-tumor brain, possibly because of rapid utilization. GMP levels, which rely on IMPDH2-mediated conversion of IMP to XMP, are increased in the tumor as compared to normal brain (Fig. 5F). We next carried out a similar experiment in vitro using ARL13B knockdown cells and steady-state metabolomics, which also demonstrated an increase in salvage pathway metabolites detected once ARl13B was lost (Sup Fig 11).

Remarkably, mice that received TMZ therapy showed decreased levels of GMP as compared to normal brain. Fractional enrichment analysis revealed that both IMP and GMP levels were higher in the tumor tissue as compared to normal brain but that TMZ therapy decreased IMP and GMP levels within the tumor significantly compared to the normal brain (Fig. 5G) Analysis of purine transporter expression revealed similar equilibrative nucleotide transporter (ENT) family levels with or without ARL13B KD (Sup. Fig. 12A).

### ARL13B knockdown increases salvage pathway mediated recycling of damaged environmental purines

Based on our *in vitro* and *in vivo* data, we propose that GBM cells downregulate salvage pathway synthesis during TMZ therapy, relying instead on de novo purine synthesis. Because TMZ derives its therapeutic efficacy by alkylating purines, we hypothesized that the inhibitory effect of ARL13B on salvage pathway might allow GBM cells to avoid recycling damaged purines from the tumor microenvironment during TMZ therapy. Moreover, GBM cells must maintain their nucleotide pools using de novo synthesis in order to continue to proliferate.

To address whether GBM cells increase de novo purine synthesis, in order to avoid utilization of TMZ-damaged purines, we first employed a modified LC-MS approach for the detection of O6-MG, the key nucleotide damage caused by TMZ treatment, in the genomic DNA of GBM cells. As shown in figure 6A, the O6-MG mark increases within 24 hrs. post-exposure to TMZ validating a positive control (m/z=166.0.723, Fig. 6A). We then analyzed ARL13B WT and KD cells treated with commercially synthesized O6-MG compound, with results showing that O6-MG mark is incorporated from the environment and recycled using salvage pathway synthesis into the DNA of ARL13B KD cells significantly more than the GBM cells with WT ARL13B (Fig. 6B: p=.025). This demonstrates that in the absence of ARL13B, a possible negative regulator of purine salvage, GBM cells salvage and incorporate damaged environmental purines into their DNA. During TMZ therapy, recycling of alkylated purines may cause extensive DNA damage, and to avoid this, GBM cells synthesize purines using the de novo pathway. This observation also supports the notion that ARL13B KD cells showed an increased propensity towards DNA damage in response to alkylating agent based chemotherapy such as TMZ and BCNU. (Fig. 6C& D, Sup. Fig. 12 B; One-way adjusted ANOVA p<.0001).

**Figure 6:**
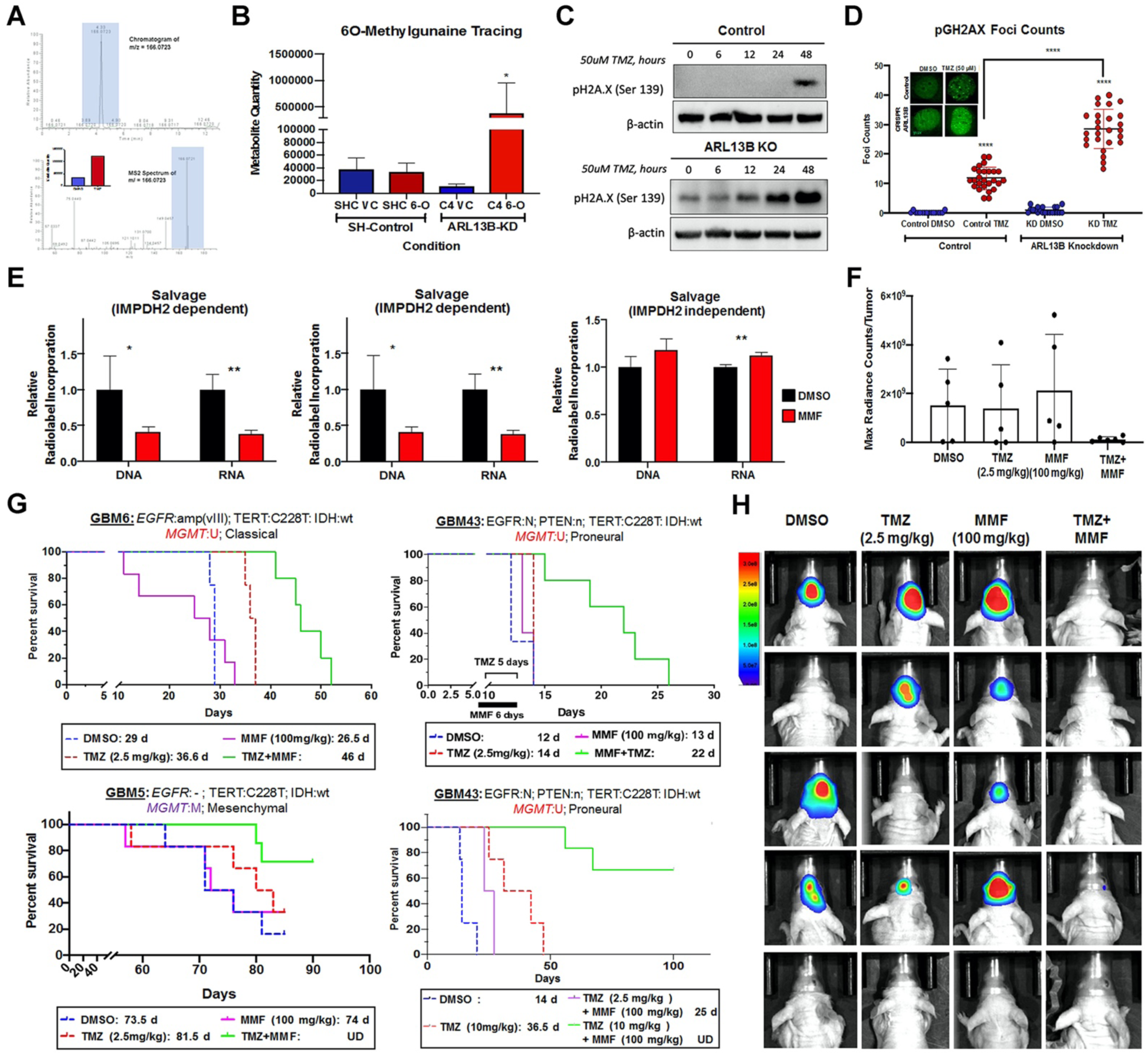
IMPDH2 inhibitor Mycophenolate Mofetil increases PDX GBM sensitivity towards TMZ *in vivo*. **A**. Chromatograms depicting peak enrichment for O6-methylguanine (O6-MG) marks in GBM cell DNA by mass spectroscopy. Graph within figure demonstrates ion count of O6-MG in cells treated with either DMSO or TMZ for 48 hours. **B**. Graph depicting O6-MG detection in samples with and without ARL13B exposed to vehicle control (VC) or O6-MG. Counts are represented as ion count with technical triplicates. **C.** Western blots analysis on cells with or without ARL13B treated with TMZ for 0, 6, 12, 24 and 48 hours and stained with pH2A.X to evaluate the extend of DNA damage. **D**: pH2A.X foci counting in cells with and without ARL13B and treated with either DMSO or TMZ (*n*=35). **E**. Radiolabeled tracing of IMPDH2-dependent salvage, de novo, and IMPDH2-independent salvage synthesis in cells exposed to DMSO control or MMF. DNA and RNA relative incorporations are shown from two biological replicates. **F.** Kaplan-Meier curves showing survival of mice implanted with classical subtypes of PDX treated with DMSO, 2.5 mg/kg TMZ for 5 days, 100mg/kg MMF for 6 days, or TMZ and MMF combination. Animals were monitored for endpoint survival. **G.** Same experiments as above but with proneural subtype GBM43, and **H.** mesenchymal subtype GBM5. **I**. Kaplan-Meier curve of mice with GBM43 PDX tumor treated with 10 mg/kg TMZ in combination with 100mg/kg MMF along with above controls **J**. BLI measurement of engrafted tumors from the mesenchymal GBM5 line followed weekly. Bar graph and images show biological replicates. Image from week three shown. All Error bars represent technical experimental replicates unless noted otherwise and display mean + SD. *p<.05 ** p<.01 ****p<.0001.

### Blocking IMPDH2 activity with Mycophenolate Moefitil sensitizes GBM cells to TMZ in vivo

Our data indicate a translational therapeutic opportunity if we can find a way to inhibit de-novo purine synthesis during TMZ therapy, forcing GBM cells to salvage alkylated purines from the environment. Unfortunately, there is no pharmacological compound that can inhibit ARL13B to replicate our phenotype. However, IMPDH2 activity can be effectively targeted by blood-brain barrier permeable compounds such as Mycophenolate Mofetil (MMF) or Mizoribine, both of which are in the clinic for transplant setting^51^. We first tested both Mizoribine as well as MMF in our mouse models and determined MMF to be better tolerated and superior to Mizoribine (Sup. Fig. 13A). MMF reduced both IMPDH2 dependent de novo purine biosynthesis (Fig. 6E; Multiple t-tests IMPDH2 Dependent Salvage: DNA p=.05, RNA p=.01, IMPDH2 Dependent de-novo: DNA p=.001, RNA p=.01, IMPDH2 independent salvage RNA p=.006). To compensate, the MMF treated GBM cells increased the IMPDH2-independent salvage pathway (Fig. 6E). Thus, cells force to use this pathway to meet their nucleotide demand and by doing so they may utilize alkylated guanosine during TMZ therapy. To validate if combining MMF with TMZ can enhance therapeutic efficacy of TMZ *in vivo*, a dose-finding experiment was done with varying MMF concentrations in combination with TMZ in which data indicated 100mg/kg to be the superior dose (Sup. Fig. 13B). Three different GBM subtypes of PDX were tested following intracranial injection of tumor cells in mice. After 7 days post-implantation animals were treated with TMZ (2.5 mg/kg), 24-hours following MMF administration (100mg/kg). In the classical GBM subtype model, this combination therapy improved median survival by 25%, 57%, and 10 % in classical, proneural, and mesenchymal models (Fig. 6F, G, & H; p =0.004, 0.01, and 0.02, respectively). The mice engrafted with GBM5, a mesenchymal line, were followed weekly with BLI to track tumor response to treatment, with results showing that MMF + TMZ is superior in slowing tumor growth (Fig. 6J). The mice from our in vivo experiment with GBM5, a mesenchymal line, were also followed weekly with BLI to show MMF + TMZ is superior in controlling the development of tumors (Fig. 6J).

*In vitro* data also suggested the MMF + TMZ combination was superior in producing DNA damage and cell killing in GBM 43 and 6, but that overexpression of ARL13B could rescue the cells from sensitivity to the MMF+ TMZ combination (Sup Fig. 14 A, B, C, D & E). To further assess the efficacy of this schema, we conducted another survival experiment with GBM43 using the same MMF dosing protocol (100mg/kg 24-hour lead-in) but now with a higher dose of TMZ (10mg/kg). This significantly enhanced the survival benefit over the TMZ alone group and may indicate that frontloading a higher TMZ dose may be more efficacious in this combination of therapy (Fig 6I TMZ alone TMZ+MMF 10mg/kg Log-Rank Adjusted p=.001).

## Discussion

Tumor cell plasticity with regards to adaptation to therapeutic pressure has been hypothesized to play a significant role in disease recurrence^12, 18, 52^. However, delineation of such adaptation mechanisms and the identification of actionable targets to prevent the plastic behavior of cancer cells are lacking. In this study, we have investigated epigenetic plasticity regulated metabolic adaptation to chemotherapy. We show that in a patient-derived xenograft model, EZH2 influences therapy-resistant BTIC frequency during alkylating-based chemotherapy by regulating ciliary protein ARL13B. We also establish that during TMZ therapy, ARL13B can interact with IMPDH2 to modulate purine biosynthesis and influence chemotherapy sensitivity. By blocking IMPDH2 activity using an FDA approved drug, we show that the therapeutic efficacy of TMZ-based therapy can be enhanced across all subtypes of GBM in PDX models.

EZH2/PRC2 is a well-studied epigenetic regulator that contributes to different malignancies, including GBM. Recent research demonstrates that BTIC cells rely on EZH2/PRC2 mediated genome-wide reposition of repressive histone marks in order to adapt to and resist targeted kinase inhibitor therapy^10^. This could also be true during TMZ therapy, and in this work, we have demonstrated that EZH2 is necessary for the frequency of BTIC’s. In order to better identify targets resulting from EZH2’s global action during therapy, we next performed an unbiased screen and identified ARL13B, as a downstream target. Our results indicated that during TMZ therapy a reduction of EZH2 binding within an ARL13B enhancer region can cause activation of ARL13B gene expression rather than silencing. However, the mechanisms of loci specific activity of EZH2/PRC2 are not known.

A key challenge with overcoming resistance is identifying an actionable target of plastic responses that are critical for tumor adaptation. As we delved into how the EZH2-ARL13B axis contributes to GBM’s response to chemotherapy, we found a novel interaction between ARL13B and IMPDH2, a critical rate-limiting enzyme for the de-novo purine biosynthesis pathway. Our data indicate that without therapeutic pressure, PDX GBM tissue maintains a high level of purine recycling salvage pathway utilization as compared to normal brain tissue. When exposed to TMZ, this heavy reliance on the salvage pathway was abolished via interaction between ARl13B and IMPDH2. This causes GBM cells switch to de-novo purine biosynthesis to meet their nucleotide demand. We hypothesized that the switch in the purine biosynthetic pathway could be due to GBM cells trying to bypass the salvage pathway in order to avoid incorporation of alkylated purines into their DNA during TMZ therapy. Our data support this notion by demonstrating when the proposed negative regulator of salvage pathway ARL13B is removed from GBM cells, and GBM cells are exposed to O6-methylguanine in their media, there is salvage mediated uptake of the alkylated purine causing it to be built into DNA. Moreover, all subtypes of GBM are more sensitive to TMZ therapy when they lack ARL13B. Pharmacologically targeting this effect by inhibiting IMPDH2 with FDA approved MMF rather than viral knockout of ARL13B also demonstrates increased efficacy of TMZ in all PDX subtypes in vivo.

Along with the current study, crucial new work has highlighted the importance of both purine biosynthesis broadly, as well as an IMPDH2 specifically, in the growth of GBM. Rich and colleagues demonstrate that the synthesis of purines utilizing the de-novo pathway is especially important in the maintenance of BTIC cells and that targeting of this can disrupt their growth potential^53^. Our results corroborate this by demonstrating reduced stem cell forming capacity when ARL13B is lost as well as the loss of expression of key stem cell markers that can ultimately be rescued by restoration of ARL13B expression. Sasaki and colleagues also show the importance of IMPDH2 and GTP biosynthesis in general for the growth of glioma^54^. They show that GBM relies on de novo GTP biosynthesis over salvage GTP biosynthesis, which dovetails with our findings that forcing salvage purine biosynthesis alone exploits a therapeutic weakness in GBM.

Our initial data points to some dependence on the nature of ARL13B as a GTPase; in fact, recent research has tied the IMP/GTP balance to the IMPDH2 function^55^. Furthermore, cells under stress in starvation conditions can form rod and ring structures termed Purinosomes^46^, which are very similar physically to cilia, possibly pointing to an avenue for a new biological process contained in cilia or the need for ARL13B to aid in forming the Purinosome itself. Moreover, it is not clear how GBM cells sense the alkylating purine in the microenvironment during TMZ therapy. Intriguingly, recent research proposed dynamic functional crosstalk between DNA damage response and cilia-associated proteins^56, 57^. Thus, it is conceivable that TMZ-induced DNA damage may influence the ciliogenesis and allow GBM cells to mount an adaptive response to promote resistance. Although our data strongly suggest the ARL13B-IMPDH2 axis can promote therapeutic adaptation in GBM, further research will be required to elucidate the precise mechanism of such resistance.

In conclusion, we present a mechanism of epigenetic plasticity that can influence purine biosynthesis pathways and allow GBM to adapt to alkylating-based chemotherapy. Our work also identifies a druggable target that can be accessed by an FDA approved compound in order to enhance the efficacy of standard of care therapy. A better understanding of these epigenetic driven metabolic adaptation processes, will be essential for developing effective therapeutic strategies against GBM.

## Acknowledgement

This work was supported by the National Institute of Neurological Disorders and Stroke grant 1R01NS096376, 1R01NS112856 the American Cancer Society grant RSG-16-034-01-DDC (to A.U.A.) grant, R01NS095642 (to C.D.J), National Institute of Cancer grant R00CA194192, National Institute of Medicine grant R01GM135587 (to I.B.-S.) and R35CA197725 (to M.S.L), P50CA221747 SPORE for Translational Approaches to Brain Cancer.

## Supplementary Figure Legends

**Supplementary Figure 1:**
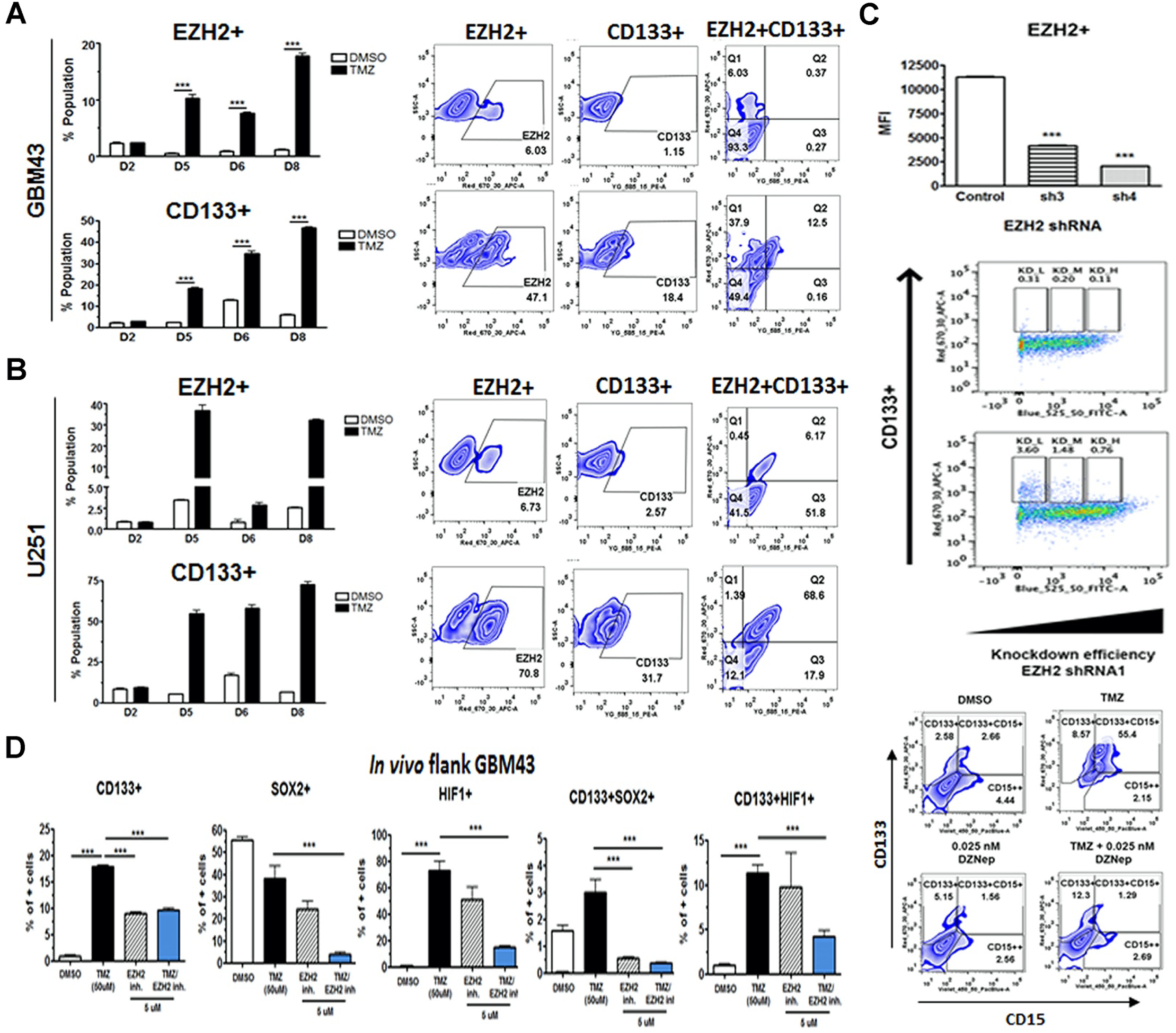
Chemical and shRNA inhibition of EZH2 impairs maintenance of BTIC niche. **A/B**: FACS plots showing percent EZH2+, CD133+, and EZH2+/CD133+ double positive populations of cells in response to TMZ therapy across two GBM cells lines. Bar graphs representing single positive populations as percent of parent (live cells). Error bars represent technical triplicates. **C**: Dot plots showing shRNA-mediated knockdown efficiency of EZH2 KD cells probed with anti-EZH2 shRNA. Bar graphs representing mean fluorescence intensity of KD cells across two different shRNA clones. **D**: FACS plots showing percent CD133+, CD15+, and CD133+/CD15+ double positive populations following treatment with DZNep with and without TMZ. Bar graphs representing single positive populations of known BTIC compartment markers as percent of parent (live cells). *** p<.001

**Supplementary Figure 2:**
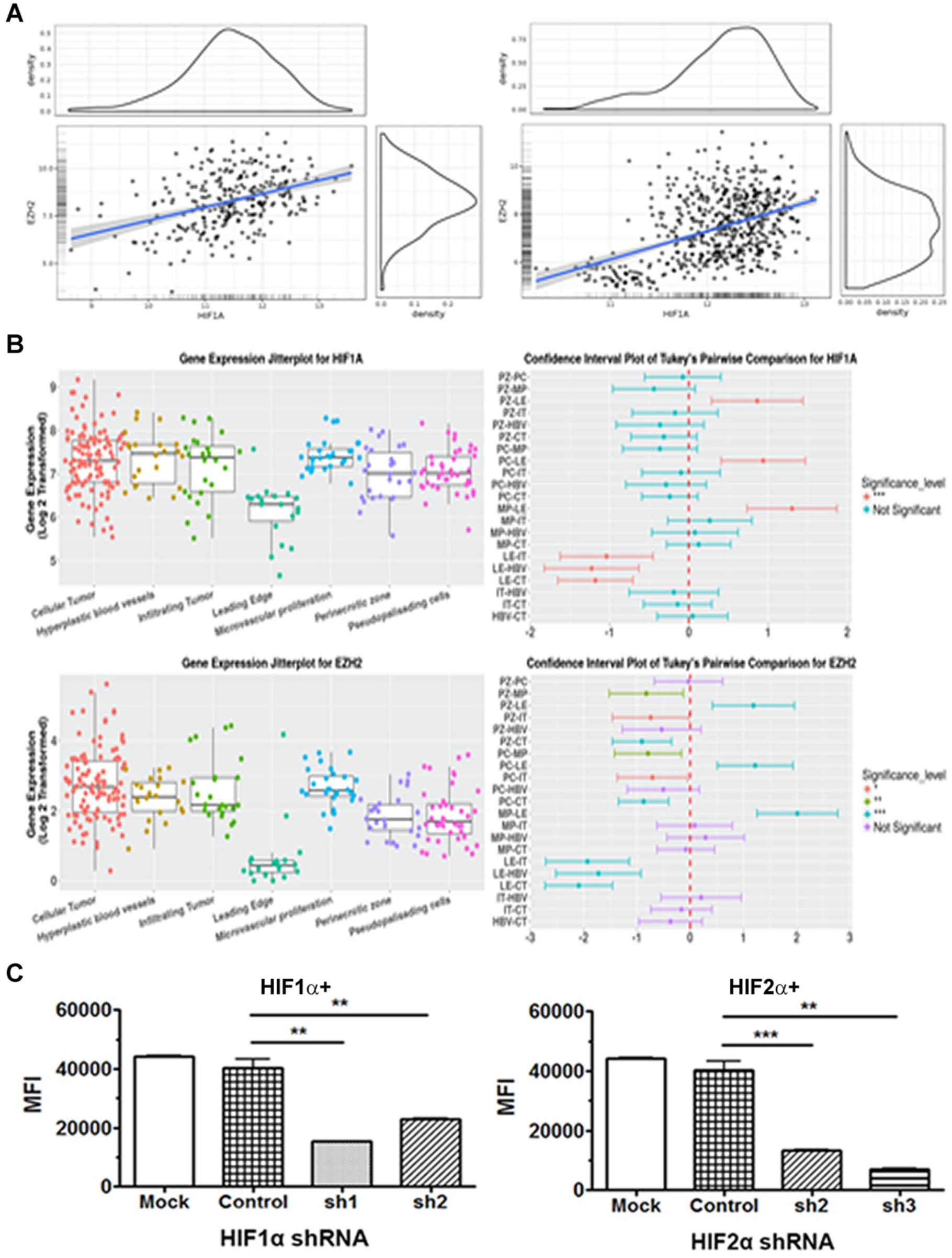
HIF1A expression is correlated with EZH2 levels in patient data. **A**: TCGA patient data showing relationship between EZH2 and HIF1A gathered from Rembrandt and Gravandeel datasets respectively. **B**: Gene expression Jitterplots showing relative transcript levels of HIF1A and EZH2 stratified by cell type. Confidence interval plots representing results of Tukey’s Pairwise Comparison for HIF1A and EZH2. **C**: Bar graphs displaying MFI of shRNA mediated KD cells stained for HIF1A and HIF2A. **p<.01 ***p<.001

**Supplementary Figure 3:**
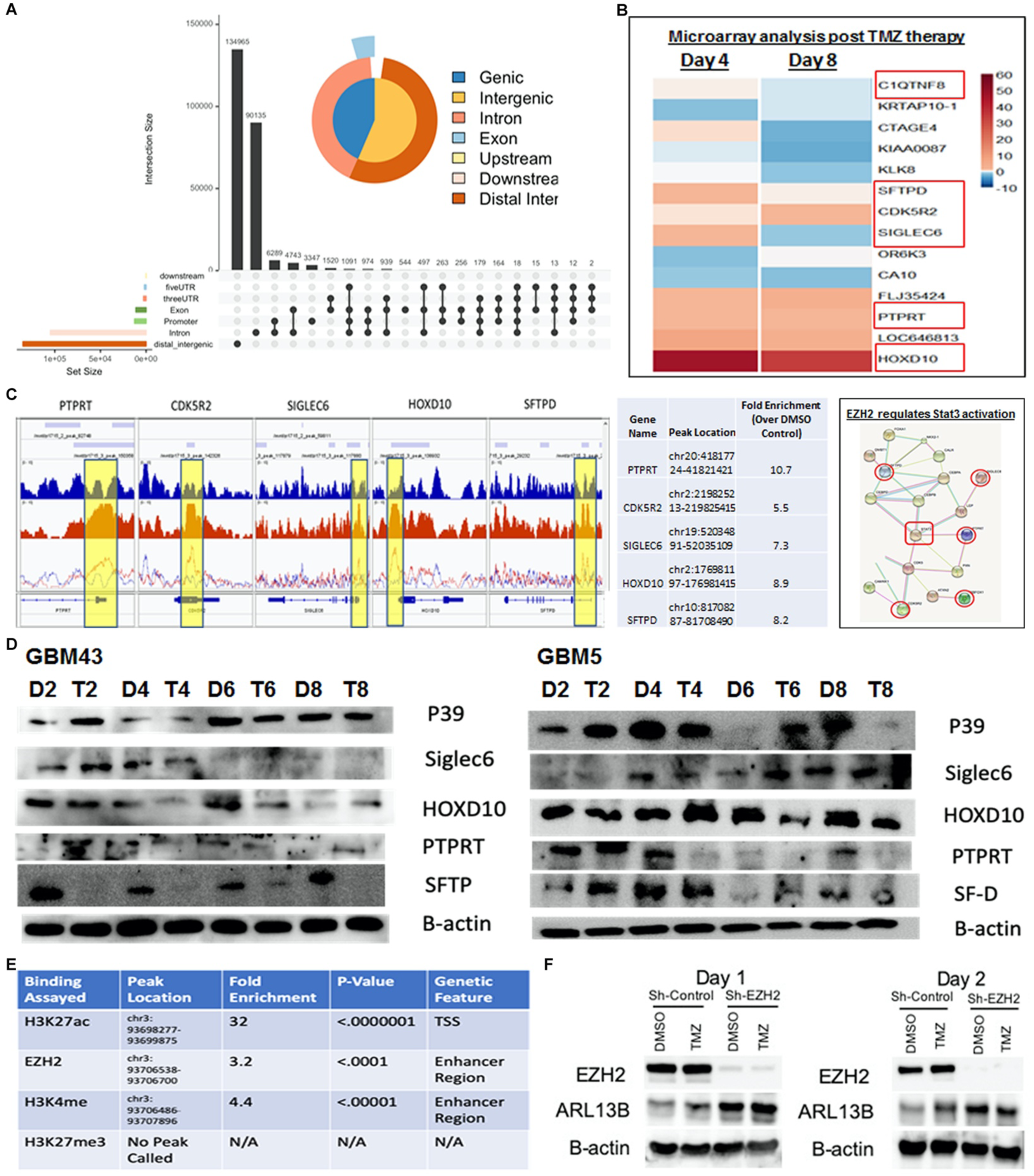
Binding of EZH2 across DNA, as well as global gene expression, is altered during TMZ therapy. **A**: Annotated VennPie plot generated from ChIPSeeker expressing distribution of EZH2 binding locations during TMZ therapy called over input DNA control. **B**: Microarray data examining global gene expression after 4 days and 8 days post TMZ therapy. **C**: ChIP data profiling H3K27ac at transcription start sites of genes found with significant alterations in expression patterns in microarray data. Yellow highlights represent statistically significant peaks post TMZ treatment that fall within the transcription start site of the genes assayed. Table indicates peak location and fold enrichment over DMSO control. String analyses of microarray targets that were also shown to have increased H3K27ac at transcription start site during TMZ therapy. **D**: Western bots profiling protein levels of these targets in GBM43 (proneural) and GBM5 (mesenchymal) cell lines. **E**: Table of multiple ChIP assays performed examining binding events within an identified ARL13B enhancer site as well as the transcription start site of ARL13B. Significance determined by BED file statistics reported from macs2 **F**: Western blot images showing expression levels of EZH2 and ARL13B under EZH2 shRNA-mediated knockdown and TMZ conditions Day 1 and 2 post therapy.

**Supplementary Figure 4:**
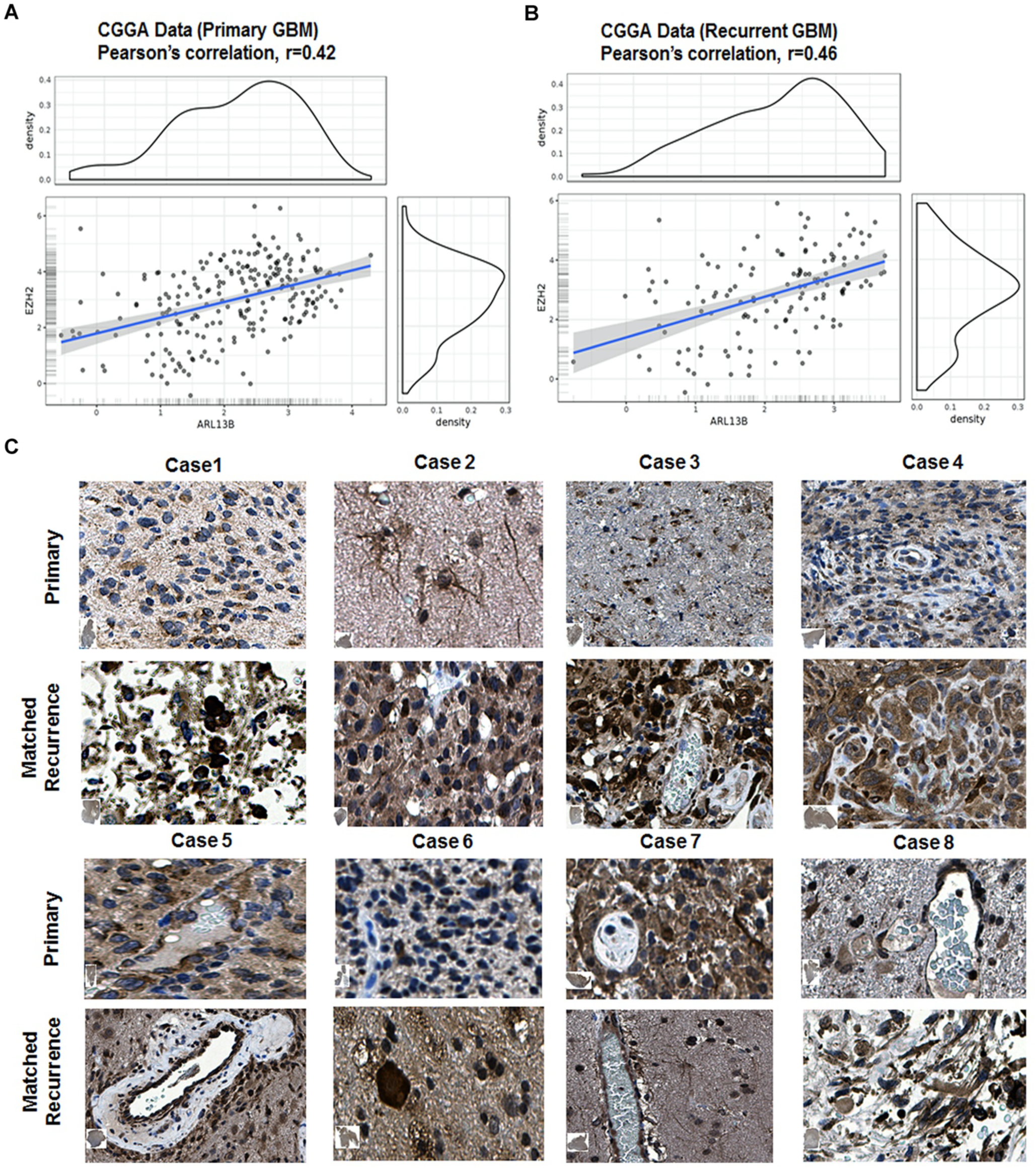
ARL13B expression is related to EZH2 levels and is correlated in both primary and recurrent tumors. **A**: CGGA data comparing expression of ARL13B and EZH2 in primary GBM **B**: CGGA data comparing expression of ARL13B and IMPDH2 in recurrent GBM. **C**: Immunohistochemistry (IHC) images of matched primary and recurrent patient samples stained for ARL13B. *n*=8.

**Supplementary Figure 5:**
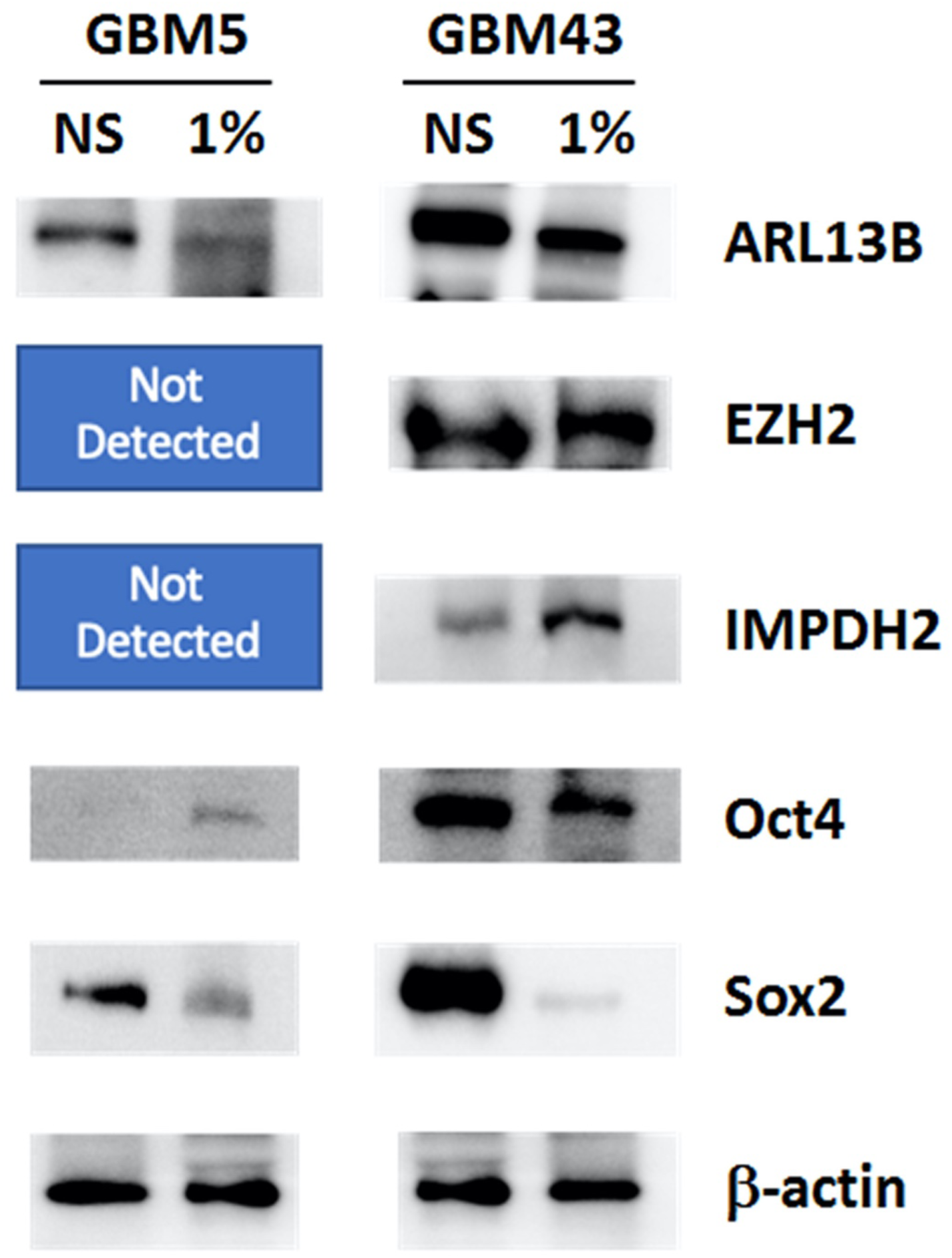
Cells cultured under stem cell conditions have increased levels of ARL13B. Western blot images comparing expression levels of several proteins in cells raised in stem cell (supplemented Neurobasil media) versus differentiated (serum grown) conditions. Data representing in vitro cultures of two different PDX lines GBM5 (mesenchymal) and GBM43 (proneural).

**Supplementary Figure 6:**
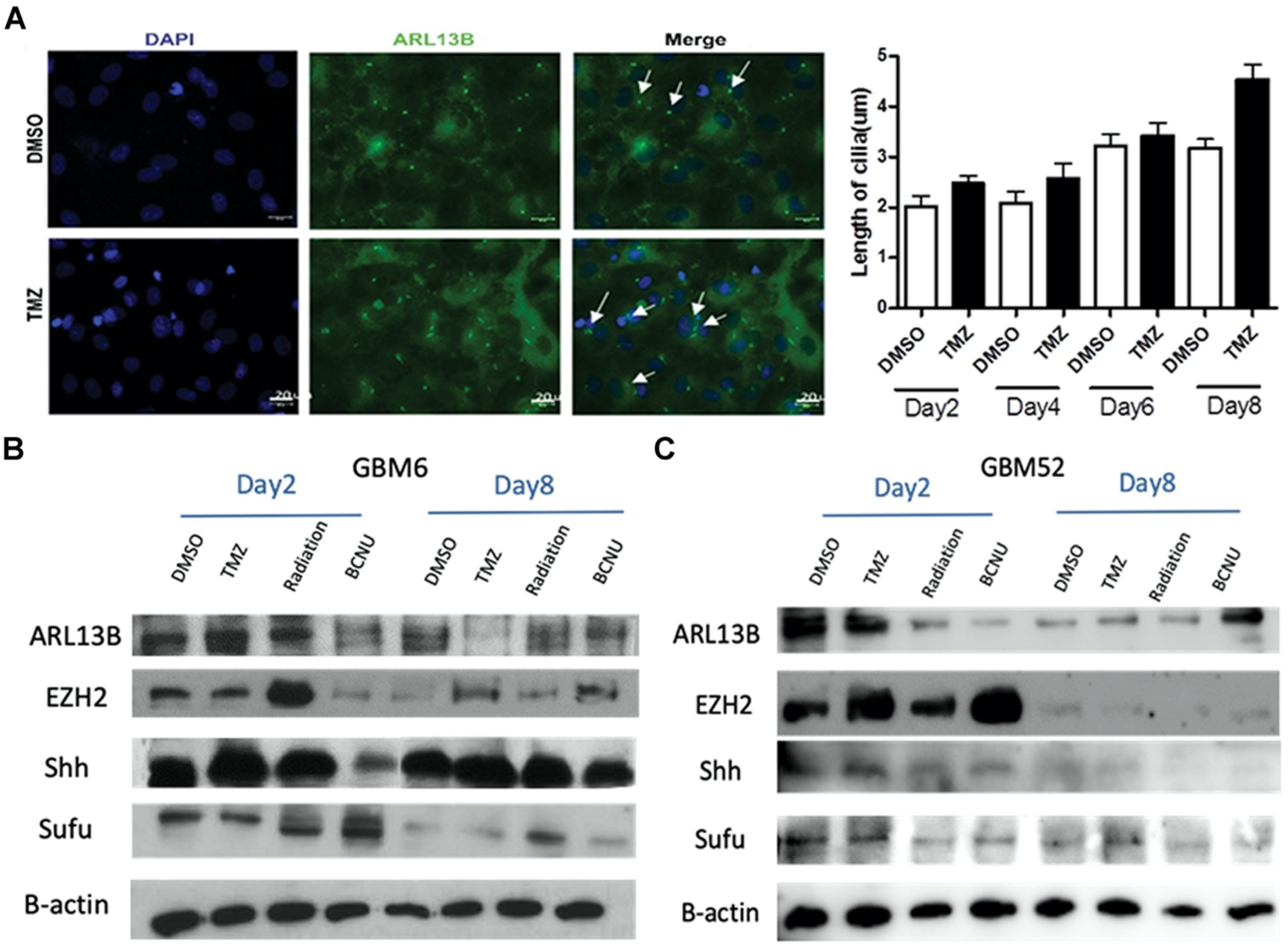
Expression of known downstream targets of ARL13B is unaltered in response to several anti-GBM treatments. **A**: Immunofluorescence (IF) images of in vitro PDX cells probed with anti-ARL13B (green) and DAPI (blue). Ciliary structures indicated by white arrows. Bar graph representing length of cilia assessed at multiple points over eight days of treatment compared to vehicle control. **B/C**: Western blot images displaying expression of various downstream targets of ARL13B following the indicated treatments across GBM6 and GBM52 cell lines.

**Supplementary Figure 7:**
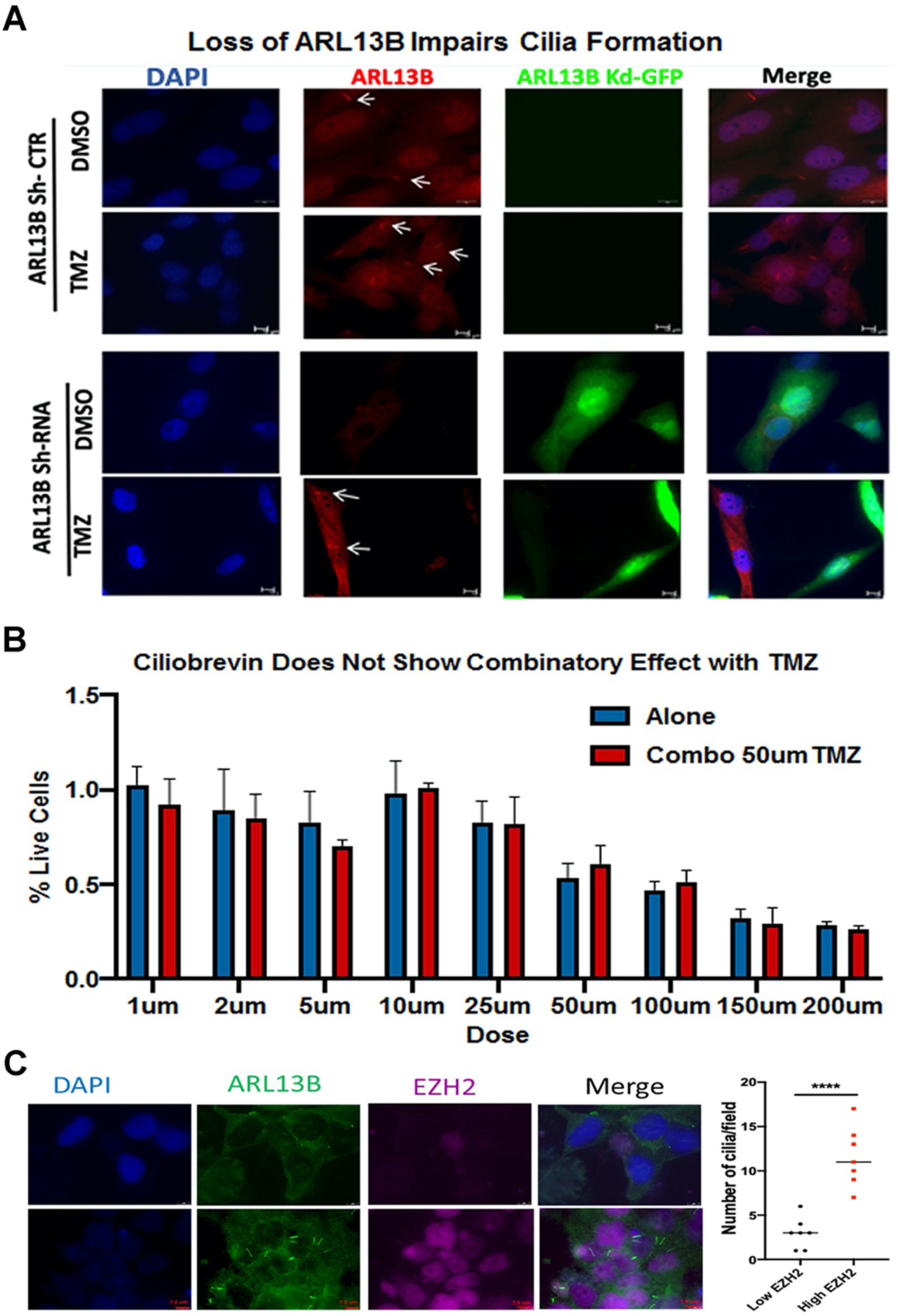
ARL13B and EZH2 expression correlate with Cilia formation but pharmacological loss of cilia does not improve response to TMZ chemotherapy. **A**: IF showing in vitro cultured cells stained with DAPI (blue), ARL13B (red), and ARL13B in knockdown condition (Green). Cells from both conditions were treated with DMSO or TMZ. **B**: IF of in vitro cultured PDX cells stained with DAPI (blue), ARL13B (green), and EZH2 (purple). Regions expressing high or low amounts of EZH2 as indicated by fluorescence intensity were isolated and quantified. Column graph visualizing number of cilia within a microscope field of view if cells have either high or low expression of EZH2 n=7 separate fields. **C**: MTT assay survival graph performed on GBM 43 pdx cells treated with varying doses of a pharmacological disruptor of cilia (Ciliobrevin) either alone or in combination with TMZ. Error bars depict technical replicates among experiments. ****p<.0001

**Supplementary Figure 8:**
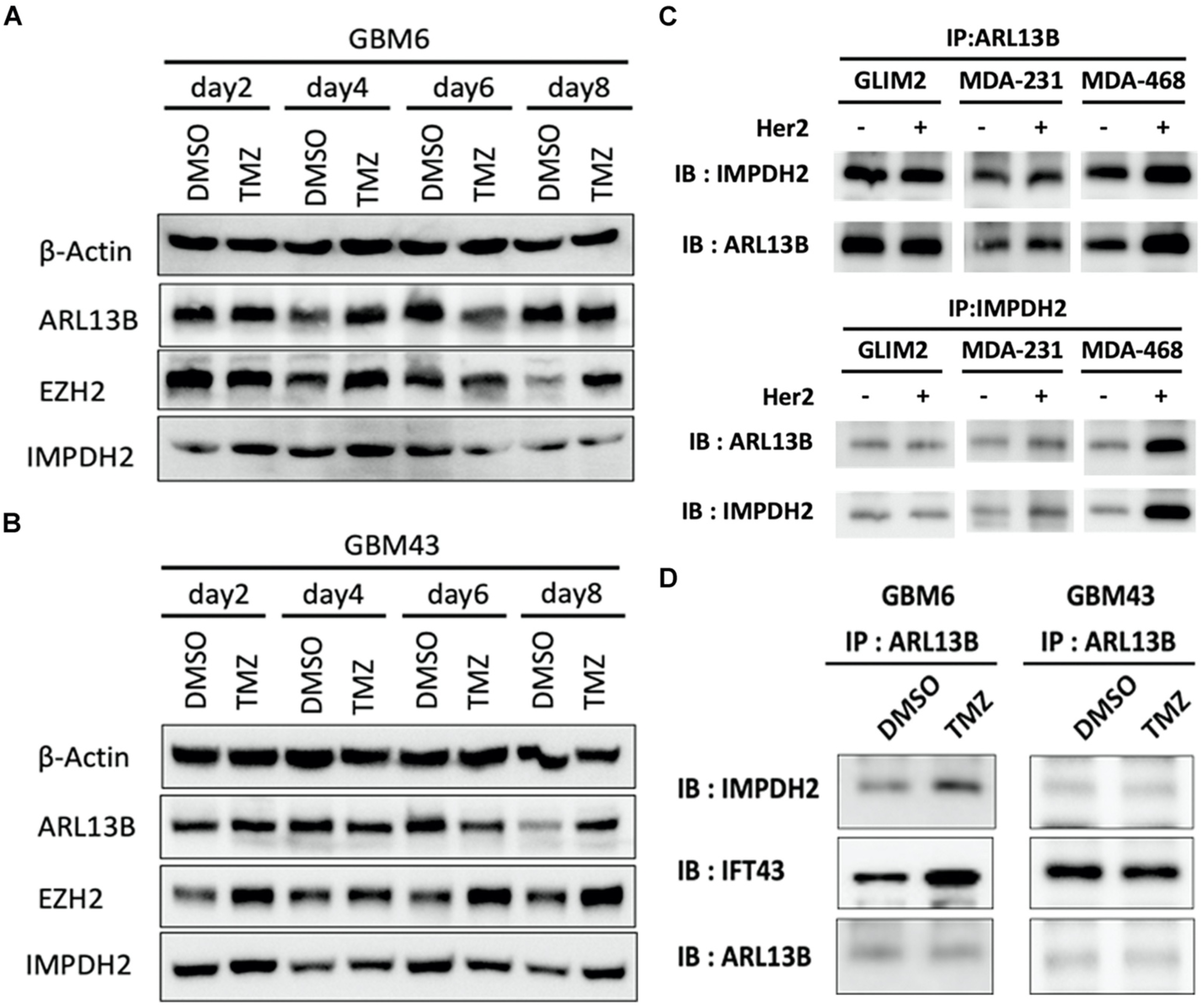
ARL13B and EZH2 protein levels increase post TMZ therapy and ARL13B and IMPDH2 interaction is also seen in breast cancer cells as well as multiple PDX subtypes. **A/B**: Western blots performed after exposure to therapy or vehicle control on GBM6 (classical) and GBM 43 (proneural). **C**: IP of ARL13B performed on cultured breast cancer cell lines probing for ARL13B and IMPDH2 interaction. **D**: IP of ARL13B performed on cultured PDX lines GBM6 (classical) and GBM43 (proneural) treated with DMSO or TMZ probing for ARL13B and IMPDH2 interaction as well as interaction with ciliary protein IF43.

**Supplementary Figure 9:**
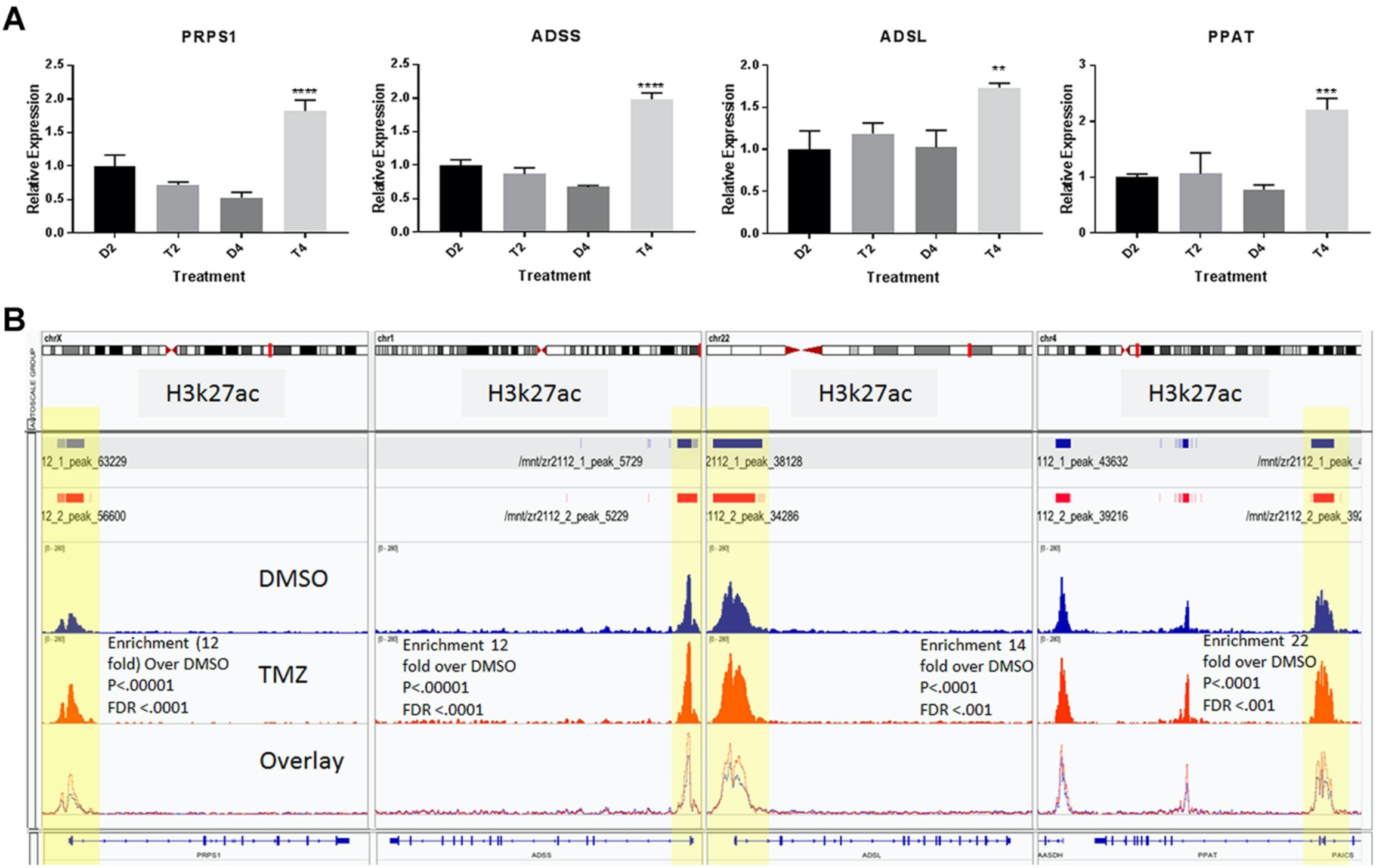
QPCR and ChipSeq demonstrate purine biosynthesis enzymes are altered post TMZ therapy. QPCR data collected on GBM 43 (proneural) across DMSO or TMZ treatment for 2 days or 4 days probing common purine biosynthetic genes. ChIPSeq demonstrated transcription start site enrichment of H3K27ac was also present. Fold enrichment and p value statistics calculated by macs2 and compared to DMSO vehicle control enrichment. Yellow highlights indicate peak within transcription start site.

**Supplementary Figure 10:**
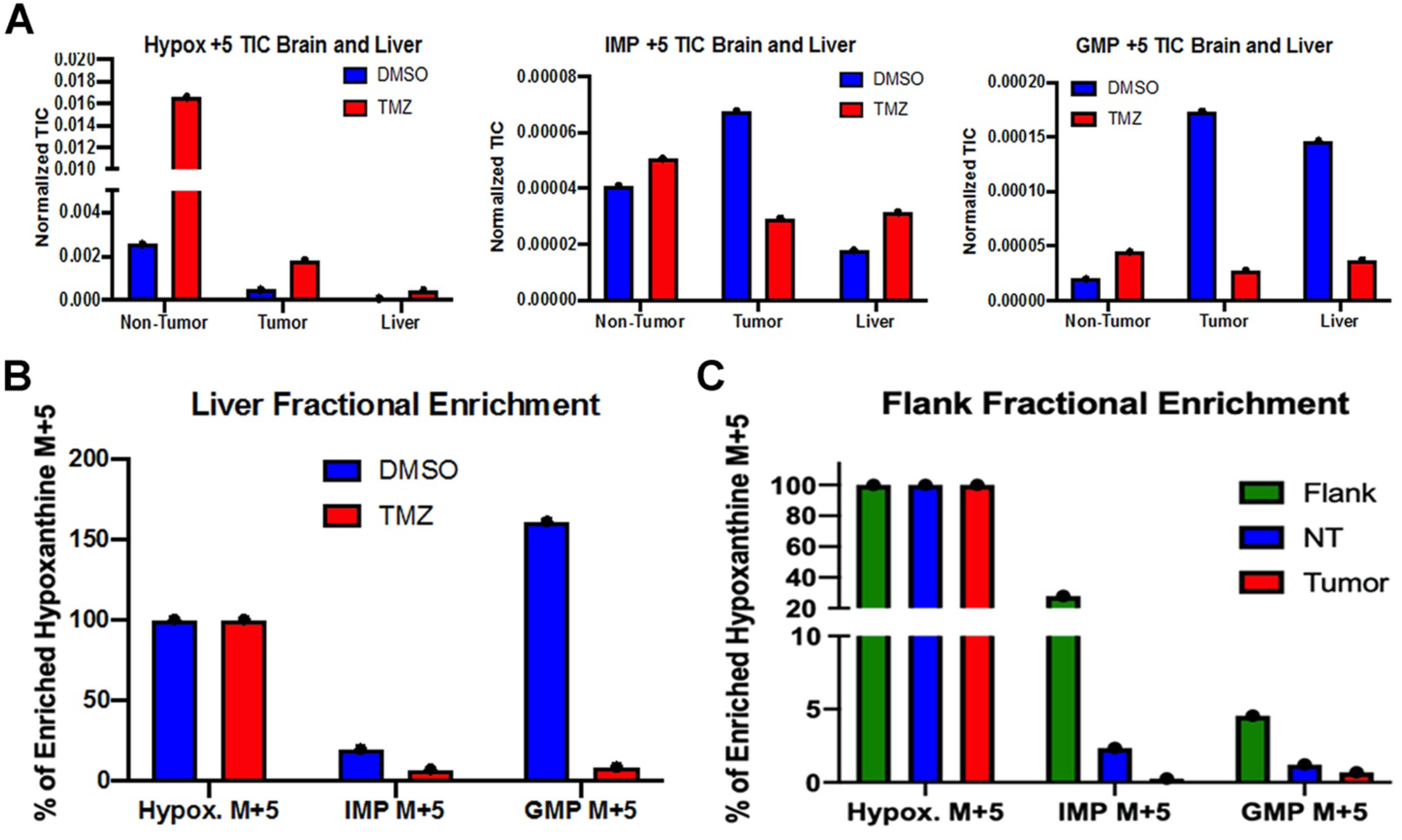
Purine turnover in the liver as well as PDX tumors grown subcutaneously in a flank show response to TMZ treatment. **A**: Graphs of total ion counts of isolated tissues from mouse brain as well as liver after mice had received either TMZ or DMSO treatment prior to hypoxanthine infusion. **B**: Fractional enrichment of IMP+5 and GMP+5 compared to Hypoxanthine +5 in the liver of mice treated with either DMSO or TMZ. **C**: Fractional enrichment of IMP+5 and GMP+5 compared to Hypoxanthine +5 in a single mouse where a GBM43 tumor was isolated from the brain as well as isolated from the subcutaneous flank and normal brain tissue.

**Supplementary Figure 11:**
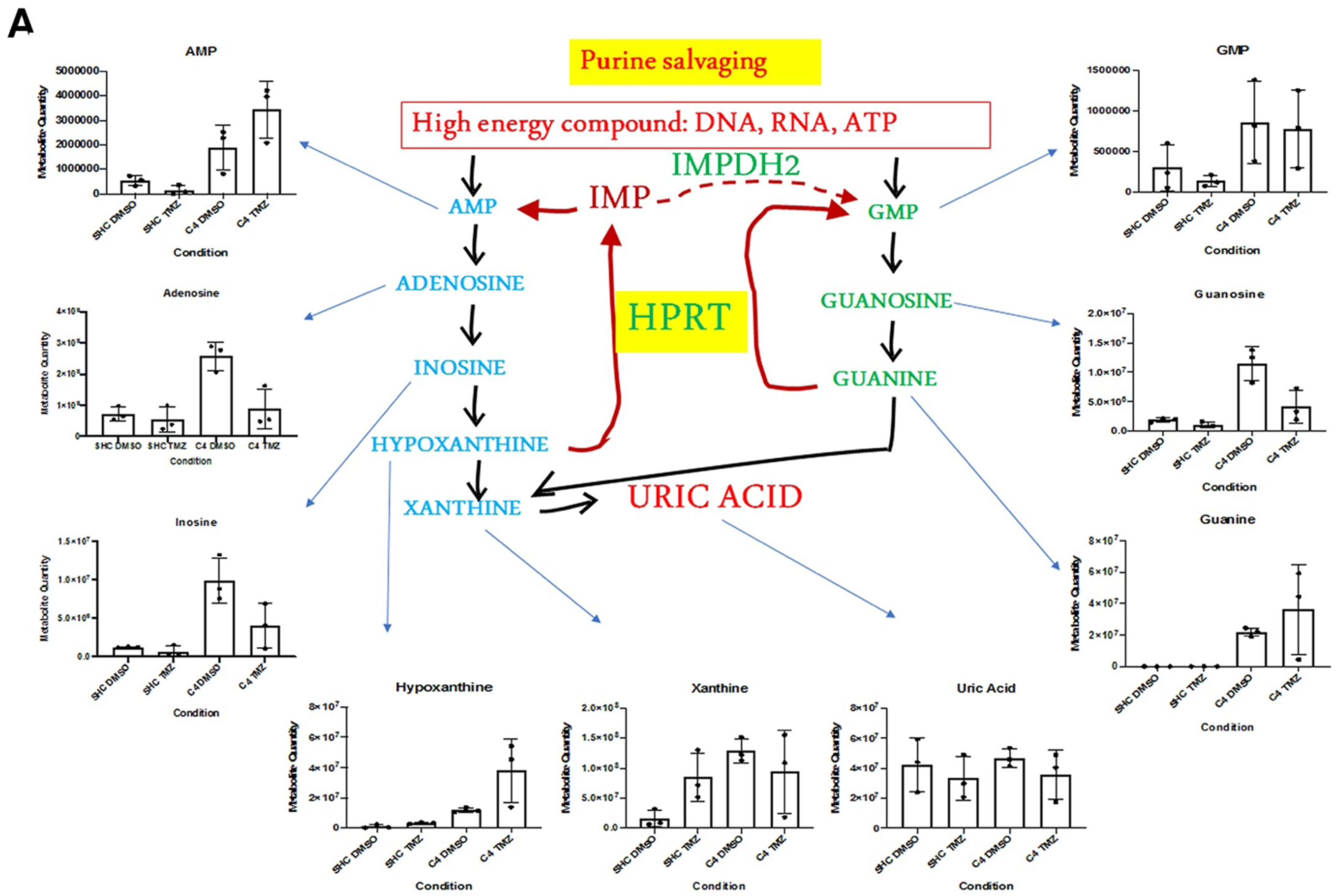
Steady state metabolic profiling of control and ARL13B knockout cells indicates altered purine biosynthesis after loss of ARl13B. Metabolites from Figure 5 in vivo tracing were assayed using cells with normal levels of ARL13B or lacking ARL13B because of CRISPR deletion. Cells were also treated with DMSO or TMZ. Error bars indicate SD of technical replicates n=3.

**Supplementary Figure 12:**
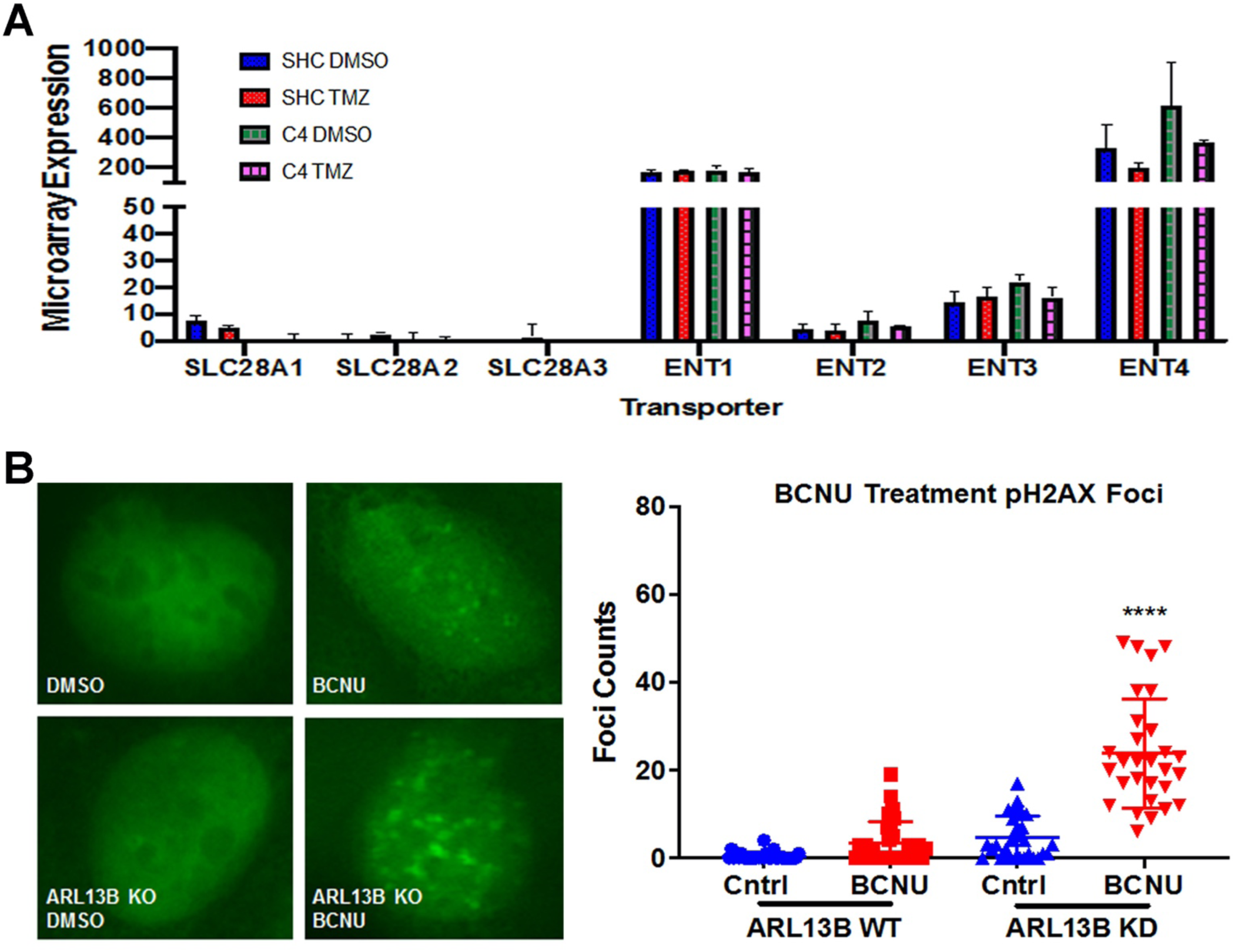
Expression of purine transporters is altered in response to TMZ and BCNU causes similar effects to TMZ with respect to DNA damge. **A**: Gene expression data from microarray analysis showing levels of several equilibrative and concentrative nucleoside transporters. **B**: Cells with or without ARL13B were treated with vehicle control or BCNU. Representative images of staining are displayed. Graph indicates blinded experimenter foci counts n=30 cells counted. **** p<.0001 2- wayANOVA adjusted for multiple comparisons.

**Supplementary Figure 13:**
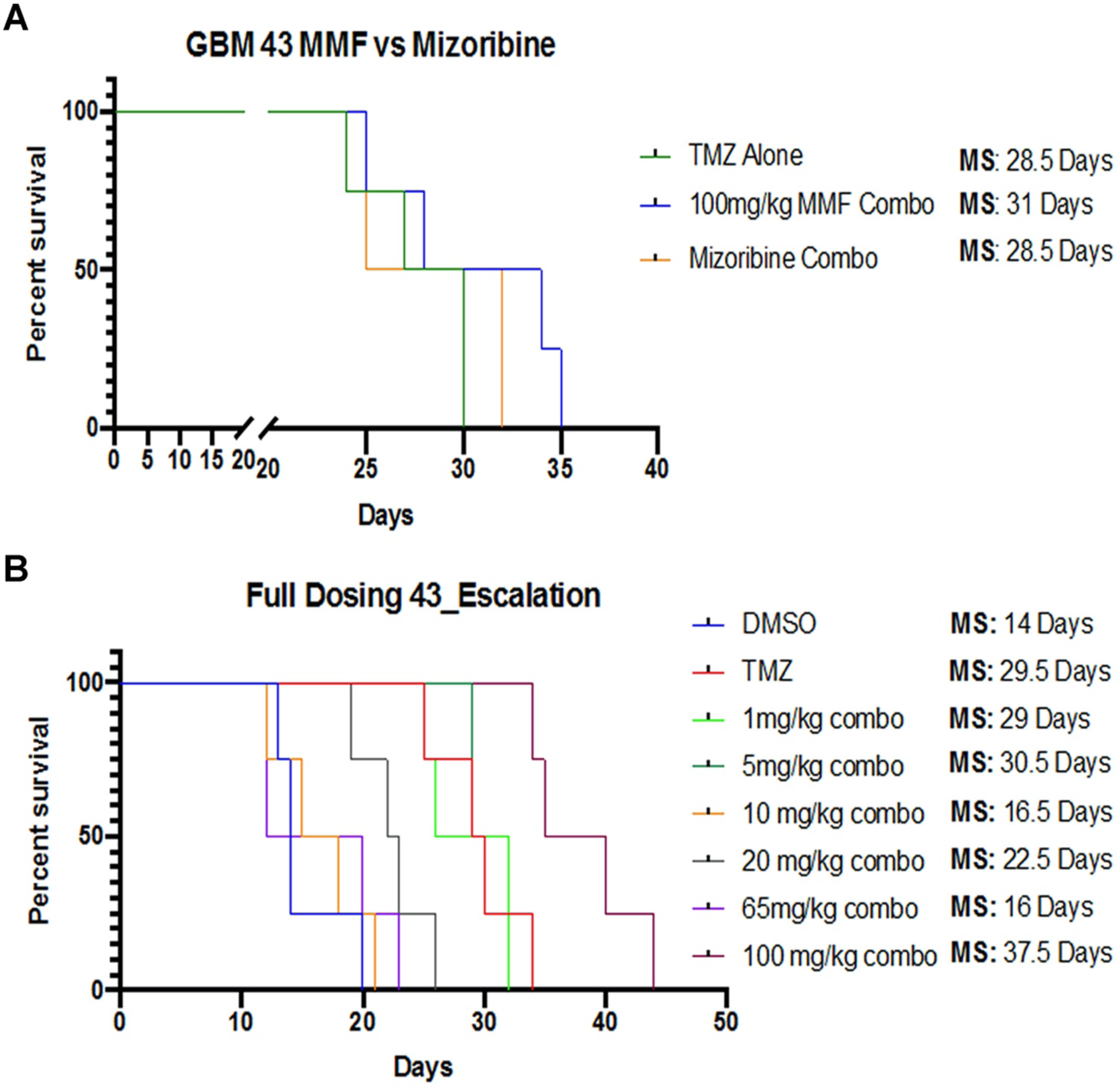
In vivo efficacy of differing doses of Mycophenolate Mofetil. **A**: Kaplan-Meier Curve representing results of experiment comparing the anti-tumor efficacy of combination MMF + TMZ versus Mizoribine+ TMZ. Median survival is indicated n=4 mice **B**: Kaplan-Meier Curve demonstrating escalating doses of MMF in combination with TMZ. Median survival is indicated n=4-5 mice.

**Supplementary Figure 14:**
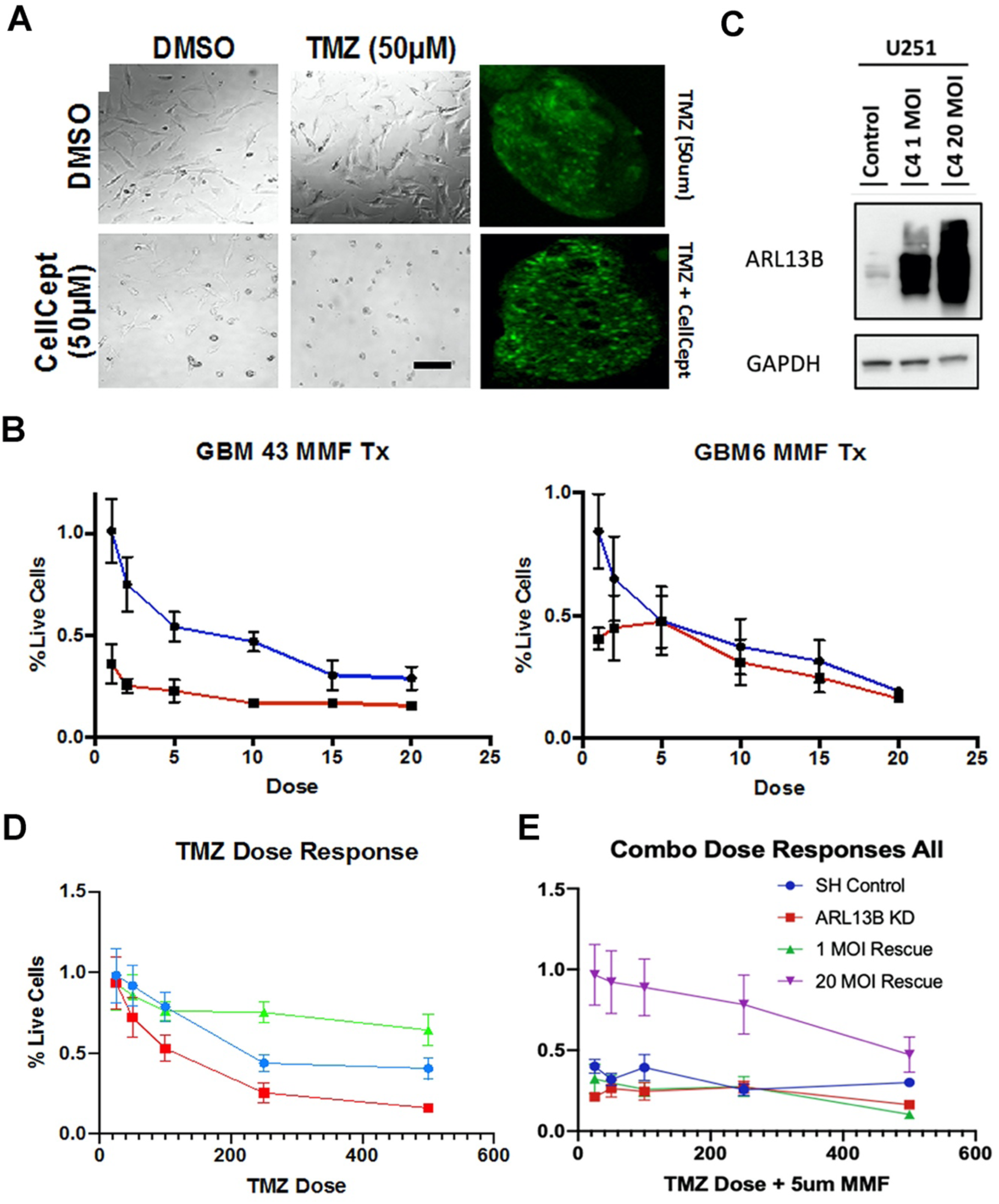
MMF causes increased DNA damage during combinatory treatment with TMZ but can be overcome with ARL13B overexpression. **A**: Representative images showing treatment with MMF (cellcept) at 50um combined with TMZ (50um) in the brightfield (live dead) and fluorescent (pH2ax) channels. **B**: In vitro survival assays done with MTT showing percent live cells in GBM 43 (proneural) and GBM6 (mesenchymal) treated with varying doses of MMF alone or in combination with 50um TMZ. **C**: Western blot in virally overexpressed U251 cells probing ARL13B protein levels across samples lacking ARL13B endogenously but infected with 1 infection unit of rescue virus or 20 infection units of rescue virus. **D**: MTT showing percent live cells treated over a TMZ dosing range from 0-500um in cells that contained normal ARL13B levels, reduced ARl13B levels, as well as rescued ARL13B overexpression. **E**: MTT assay showing percent live cells across TMZ treatments ranging from 0-500um combined with 5um MMF in cells with WT ARL13B expression, Loss of ARL13B, rescue of ARL13B expression, and rescue overexpression of ARL13B.

## References

1. Fabian, D., et al. Treatment of Glioblastoma (GBM) with the Addition of Tumor-Treating Fields (TTF): A Review. Cancers (Basel) 11(2019).

2. Stupp, R. & Ram, Z. Quality of Life in Patients With Glioblastoma Treated With Tumor-Treating Fields-Reply. JAMA 319, 1823 (2018).

3. Fidoamore, A., et al. Glioblastoma Stem Cells Microenvironment: The Paracrine Roles of the Niche in Drug and Radioresistance. Stem Cells Int 2016, 6809105 (2016).

4. Sharifzad, F., et al. Glioblastoma cancer stem cell biology: Potential theranostic targets. Drug Resist Updat 42, 35–45 (2019).

5. Bernstock, J.D., et al. Molecular and cellular intratumoral heterogeneity in primary glioblastoma: clinical and translational implications. J Neurosurg, 1–9 (2019).

6. Heddleston, J.M., Li, Z., McLendon, R.E., Hjelmeland, A.B. & Rich, J.N. The hypoxic microenvironment maintains glioblastoma stem cells and promotes reprogramming towards a cancer stem cell phenotype. Cell Cycle 8, 3274–3284 (2009).

7. Chaffer, C.L., et al. Normal and neoplastic nonstem cells can spontaneously convert to a stem-like state. Proceedings of the National Academy of Sciences 108, 7950 (2011).

8. Auffinger, B., et al. Conversion of differentiated cancer cells into cancer stem-like cells in a glioblastoma model after primary chemotherapy. Cell Death Differ 21, 1119–1131 (2014).

9. Dahan, P., et al. Ionizing radiations sustain glioblastoma cell dedifferentiation to a stem-like phenotype through survivin: possible involvement in radioresistance. Cell Death Dis 5, e1543 (2014).

10. Liau, B.B., et al. Adaptive Chromatin Remodeling Drives Glioblastoma Stem Cell Plasticity and Drug Tolerance. Cell Stem Cell 20, 233–246 e237 (2017).

11. Safa, A.R., Saadatzadeh, M.R., Cohen-Gadol, A.A., Pollok, K.E. & Bijangi-Vishehsaraei, K. Glioblastoma stem cells (GSCs) epigenetic plasticity and interconversion between differentiated non-GSCs and GSCs. Genes Dis 2, 152–163 (2015).

12. Sheahan, A.V. & Ellis, L. Epigenetic reprogramming: A key mechanism driving therapeutic resistance. Urol Oncol 36, 375–379 (2018).

13. Qi, L., et al. The dynamics of polycomb group proteins in early embryonic nervous system in mouse and human. Int J Dev Neurosci 31, 487–495 (2013).

14. Zhao, L., et al. Ezh2 is involved in radial neuronal migration through regulating Reelin expression in cerebral cortex. Sci Rep 5, 15484 (2015).

15. Deevy, O. & Bracken, A.P. PRC2 functions in development and congenital disorders. Development 146(2019).

16. Suva, M.L., et al. EZH2 Is Essential for Glioblastoma Cancer Stem Cell Maintenance. Cancer Research 69, 9211–9218 (2009).

17. Orzan, F., et al. Enhancer of Zeste 2 (EZH2) is up-regulated in malignant gliomas and in glioma stem-like cells. Neuropathol Appl Neurobiol 37, 381–394 (2011).

18. Natsume, A., et al. Chromatin regulator PRC2 is a key regulator of epigenetic plasticity in glioblastoma. Cancer Res 73, 4559–4570 (2013).

19. Kim, E., et al. Phosphorylation of EZH2 activates STAT3 signaling via STAT3 methylation and promotes tumorigenicity of glioblastoma stem-like cells. Cancer Cell 23, 839–852 (2013).

20. Wang, P., et al. HIF1alpha regulates glioma chemosensitivity through the transformation between differentiation and dedifferentiation in various oxygen levels. Sci Rep 7, 7965 (2017).

21. Caragher, S.P., et al. Activation of Dopamine Receptor 2 Prompts Transcriptomic and Metabolic Plasticity in Glioblastoma. The Journal of Neuroscience 39, 1982 (2019).

22. Oizel, K., et al. Efficient Mitochondrial Glutamine Targeting Prevails Over Glioblastoma Metabolic Plasticity. Clin Cancer Res 23, 6292–6304 (2017).

23. Sarkaria, J.N., et al. Use of an orthotopic xenograft model for assessing the effect of epidermal growth factor receptor amplification on glioblastoma radiation response. Clin Cancer Res 12, 2264–2271 (2006).

24. Yuan, M., Breitkopf, S.B., Yang, X. & Asara, J.M. A positive/negative ion-switching, targeted mass spectrometry-based metabolomics platform for bodily fluids, cells, and fresh and fixed tissue. Nat Protoc 7, 872–881 (2012).

25. Ramirez, F., et al. High-resolution TADs reveal DNA sequences underlying genome organization in flies. Nat Commun 9, 189 (2018).

26. Ben-Sahra, I., Howell, J.J., Asara, J.M. & Manning, B.D. Stimulation of de novo pyrimidine synthesis by growth signaling through mTOR and S6K1. Science 339, 1323–1328 (2013).

27. Beier, D., et al. Temozolomide preferentially depletes cancer stem cells in glioblastoma. Cancer Res 68, 5706–5715 (2008).

28. Rosso, L., et al. A new model for prediction of drug distribution in tumor and normal tissues: pharmacokinetics of temozolomide in glioma patients. Cancer Res 69, 120–127 (2009).

29. Ostermann, S., et al. Plasma and cerebrospinal fluid population pharmacokinetics of temozolomide in malignant glioma patients. Clin Cancer Res 10, 3728–3736 (2004).

30. Li, Z., et al. Hypoxia-inducible factors regulate tumorigenic capacity of glioma stem cells. Cancer Cell 15, 501–513 (2009).

31. Chang, C.J., et al. EZH2 promotes expansion of breast tumor initiating cells through activation of RAF1-beta-catenin signaling. Cancer Cell 19, 86–100 (2011).

32. Lee, G., et al. Dedifferentiation of Glioma Cells to Glioma Stem-like Cells By Therapeutic Stress-induced HIF Signaling in the Recurrent GBM Model. Mol Cancer Ther 15, 3064–3076 (2016).

33. Bowman, R.L., Wang, Q., Carro, A., Verhaak, R.G. & Squatrito, M. GlioVis data portal for visualization and analysis of brain tumor expression datasets. Neuro Oncol 19, 139–141 (2017).

34. Chang, N., Ahn, S.H., Kong, D.S., Lee, H.W. & Nam, D.H. The role of STAT3 in glioblastoma progression through dual influences on tumor cells and the immune microenvironment. Mol Cell Endocrinol 451, 53–65 (2017).

35. Zhang, J., et al. EZH2 is a negative prognostic factor and exhibits pro-oncogenic activity in glioblastoma. Cancer Lett 356, 929–936 (2015).

36. Gao, T., et al. EnhancerAtlas: a resource for enhancer annotation and analysis in 105 human cell/tissue types. Bioinformatics 32, 3543–3551 (2016).

37. Lee, J., et al. Tumor stem cells derived from glioblastomas cultured in bFGF and EGF more closely mirror the phenotype and genotype of primary tumors than do serum-cultured cell lines. Cancer Cell 9, 391–403 (2006).

38. Tobias, A.L., et al. The Timing of Neural Stem Cell-Based Virotherapy Is Critical for Optimal Therapeutic Efficacy When Applied With Radiation and Chemotherapy for the Treatment of Glioblastoma. 2, 655–666 (2013).

39. Jeng, K.S., Chang, C.F. & Lin, S.S. Sonic Hedgehog Signaling in Organogenesis, Tumors, and Tumor Microenvironments. Int J Mol Sci 21(2020).

40. Bay, S.N., Long, A.B. & Caspary, T. Disruption of the ciliary GTPase Arl13b suppresses Sonic hedgehog overactivation and inhibits medulloblastoma formation. Proc Natl Acad Sci U S A 115, 1570–1575 (2018).

41. Mariani, L.E., et al. Arl13b regulates Shh signaling from both inside and outside the cilium. Molecular Biology of the Cell 27, 3780–3790 (2016).

42. Larkins, C.E., Aviles, G.D., East, M.P., Kahn, R.A. & Caspary, T. Arl13b regulates ciliogenesis and the dynamic localization of Shh signaling proteins. Mol Biol Cell 22, 4694–4703 (2011).

43. Gajjar, A.J. & Robinson, G.W. Medulloblastoma—translating discoveries from the bench to the bedside. Nature Reviews Clinical Oncology 11, 714–722 (2014).

44. Camici, M., Garcia-Gil, M., Pesi, R., Allegrini, S. & Tozzi, M.G. Purine-Metabolising Enzymes and Apoptosis in Cancer. Cancers (Basel) 11(2019).

45. Hedstrom, L. IMP dehydrogenase: structure, mechanism, and inhibition. Chem Rev 109, 2903–2928 (2009).

46. Pedley, A.M. & Benkovic, S.J. A New View into the Regulation of Purine Metabolism: The Purinosome. Trends Biochem Sci 42, 141–154 (2017).

47. Ipata, P.L., Camici, M., Micheli, V. & Tozz, M.G. Metabolic network of nucleosides in the brain. Curr Top Med Chem 11, 909–922 (2011).

48. Micheli, V., et al. Neurological disorders of purine and pyrimidine metabolism. Curr Top Med Chem 11, 923–947 (2011).

49. Seegmiller, J.E., Rosenbloom, F.M. & Kelley, W.N. Enzyme Defect Associated with a Sex-Linked Human Neurological Disorder and Excessive Purine Synthesis. Science 155, 1682–1684 (1967).

50. Brada, M., et al. Phase I dose-escalation and pharmacokinetic study of temozolomide (SCH 52365) for refractory or relapsing malignancies. Br J Cancer 81, 1022–1030 (1999).

51. Allison, A.C. & Eugui, E.M. Purine metabolism and immunosuppressive effects of mycophenolate mofetil (MMF). Clinical transplantation 10, 77–84 (1996).

52. Das, P.K., et al. Plasticity of Cancer Stem Cell: Origin and Role in Disease Progression and Therapy Resistance. Stem Cell Rev Rep (2020).

53. Wang, X., et al. Purine synthesis promotes maintenance of brain tumor initiating cells in glioma. Nat Neurosci 20, 661–673 (2017).

54. Kofuji, S., et al. IMP dehydrogenase-2 drives aberrant nucleolar activity and promotes tumorigenesis in glioblastoma. Nat Cell Biol 21, 1003–1014 (2019).

55. Keppeke, G.D., et al. IMP/GTP balance modulates cytoophidium assembly and IMPDH activity. Cell Div 13, 5 (2018).

56. Villumsen, B.H., et al. A new cellular stress response that triggers centriolar satellite reorganization and ciliogenesis. EMBO J 32, 3029–3040 (2013).

57. Johnson, C.A. & Collis, S.J. Ciliogenesis and the DNA damage response: a stressful relationship. Cilia 5, 19 (2016).

